# Talin and vinculin combine their activities to trigger actin assembly

**DOI:** 10.1101/2022.12.01.518757

**Authors:** Hong Wang, Clémence Vigouroux, Rayan Said, Véronique Henriot, Julien Pernier, Christophe Le Clainche

## Abstract

Focal adhesions (FAs) strengthen their link with the actin cytoskeleton to resist force. Talin-vinculin association could reinforce actin anchoring to FAs by controlling actin polymerization. However, the actin polymerization activity of the talin-vinculin complex is not known because it requires the reconstitution of the mechanical and biochemical activation steps that control the association of talin and vinculin and the exposure of their actin-binding domains. By combining kinetic and binding assays with single actin filament observations in TIRF microscopy, we show that the association of talin and vinculin mutants, mimicking different degrees of activation, results in a variety of activities. In particular, mechanically stretched talin and activated vinculin combine to stimulate actin assembly synergistically through a sequential mechanism in which filaments are nucleated, capped and released to elongate. Our findings suggest a versatile mechanism for the regulation of actin assembly in FAs subjected to various combinations of biochemical and mechanical cues.

## Introduction

To migrate efficiently in different tissues, cells must sense and adapt to variations of the mechanical properties of their environment. In this adaptive process, focal adhesions (FAs) can strengthen their link with the extracellular matrix (ECM) and the actomyosin stress fibers (Ciobanasu et al., 2013; Gardel et al., 2010; Geiger et al., 2009). FAs are composed of transmembrane integrins that mechanically couple the ECM to the actomyosin cytoskeleton, via a variety of actin binding proteins (ABPs) (Bachmann et al., 2019; Ciobanasu et al., 2013; Le Clainche and Carlier, 2008; Romero et al., 2020).

The mechanical coupling of stress fibers to FAs is highly regulated. The anchoring of actin filaments to FAs can be modulated by the degree of engagement of a molecular clutch composed of sliding layers of interacting proteins, including ABPs (Gardel et al., 2010; Hu et al., 2007). The regulation of the polymerization of the actin filaments that compose the stress fibers may also determine the level of their mechanical coupling to FAs (Ciobanasu et al., 2013). Interestingly, FAs associated with elongating dorsal stress fibers are associated with low traction forces, whereas FAs associated with non-elongating ventral stress fibers are associated with high traction forces (Tojkander et al., 2015). This inverse correlation between the elongation of the actin filaments and the transmission of force to the ECM sheds light on the importance of force-dependent ABPs which control actin assembly in FAs.

Biochemical and cellular studies have described a variety of ABPs associated with FAs that regulate the elongation of actin filament barbed ends. Early in vitro studies showed that the vasodilator-stimulated phosphoprotein (VASP) nucleates actin filaments and assembles them into bundles (Laurent et al., 1999). More recent studies demonstrate that VASP can also elongate actin filament barbed ends in a processive manner (Breitsprecher et al., 2011, 2008; Hansen and Mullins, 2010). Similarly, a series of studies suggested that formins are involved in the processive elongation of stress fibers in cells (Hotulainen and Lappalainen, 2006; Tojkander et al., 2011). Talin and vinculin, which associate in response to the actomyosin force, could link force sensing to the control of actin polymerization in FAs.

Vinculin is an autoinhibited ABP in which the single actin-binding domain (ABD), called vinculin tail (V_t_), is masked by an intramolecular interaction with vinculin head (V_h_) (Johnson and Craig, 1995). Biochemical studies showed that isolated V_t_ binds actin filaments and assembles them into bundles (Huttelmaier et al., 1997; Janssen et al., 2006). We showed that V_t_ also caps actin filament barbed ends and nucleates actin filaments (Le Clainche et al., 2010). All these activities of V_t_ are masked by V_h_ in the autoinhibited conformation of vinculin.

Talin is a large ABP composed of a FERM domain, subdivided into F0, F1, F2, F3, and a rod domain made of 13 helical bundles (R1 to R13) (Ciobanasu et al., 2013). The two major intramolecular interactions F3-R9 and F2-R12 keep talin in an autoinhibited form (Dedden et al., 2019; Goult et al., 2009). Talin contains three ABDs. ABD1, ABD2 and ABD3 correspond to the head (F2F3), the central region (R4-R8) and the last C-terminal bundle (R13) respectively (Hemmings et al., 1996). We previously demonstrated that the N-terminal ABD1 of talin blocks the elongation of actin filament barbed ends in low ionic strength conditions, whereas ABD2 and ABD3 do not affect actin dynamics (Ciobanasu et al., 2018). Full-length talin is inactive because ABD1 is inhibited by the F3-R9 intramolecular interaction.

Biochemical and structural studies revealed the presence of 11 vinculin binding sites (VBSs) buried in some of the 13 helical bundles of talin (Gingras et al., 2005). The stretching of single molecules of talin revealed that force exposes several of these 11 cryptic VBSs along talin rod domain (del Rio et al., 2009; Goult et al., 2018; Yao et al., 2016). Using an in vitro reconstitution approach, we demonstrated that the actomyosin force is sufficient to stretch talin, allowing its binding to vinculin (Ciobanasu et al., 2014). The release of talin autoinhibition also favors the low-affinity constitutive binding of vinculin (Atherton et al., 2020).

In order for the V_h_ domain of vinculin to interact with talin VBSs, it is necessary to release it from its autoinhibitory interaction with V_t_ (Johnson and Craig, 1994). Although the mechanism of activation of vinculin is not fully understood, several studies show that once vinculin is bound to talin, the interaction of V_t_ with actomyosin keeps vinculin under tension in its open conformation (Carisey et al., 2013; Chen et al., 2006; Grashoff et al., 2010; Hirata et al., 2014; Humphries et al., 2007). In this mechanism, the actomyosin-dependent talin-vinculin complex acts as a catch-bond that releases V_t_ to strengthen the anchoring to actomyosin whose contraction initially induced its formation (Atherton et al., 2015; Ciobanasu et al., 2014; Hirata et al., 2014).

Because force is required to trigger vinculin association to talin, and because the two proteins are autoinhibited, it has so far been impossible to determine the ability of the talin-vinculin complex to regulate actin polymerization. Here, we designed a series of talin and vinculin mutants that associate constitutively into a stable complex. By combining kinetic studies in fluorescence spectroscopy, actin binding assays and single actin filament observations in TIRF microscopy, we determined the activities of these mutants and their complexes on actin binding and polymerization. Altogether our data reveal a force-dependent mechanism of actin assembly in FAs.

## Results

### The release of the two autoinhibitory contacts of vinculin allows F-actin binding and barbed-end capping but not nucleation

Before starting the study of the talin-vinculin complex, we deciphered the complex mechanism that links the autoinhibition of vinculin and its activities. Indeed, biochemical and structural studies have revealed that the auto-inhibition of vinculin is controlled by two interfaces between V_t_ and the D1 and D4 subdomains of the head (Bakolitsa et al., 2004; Cohen et al., 2005; Le Clainche et al., 2010) (Figure 1A).

**Figure 1.**
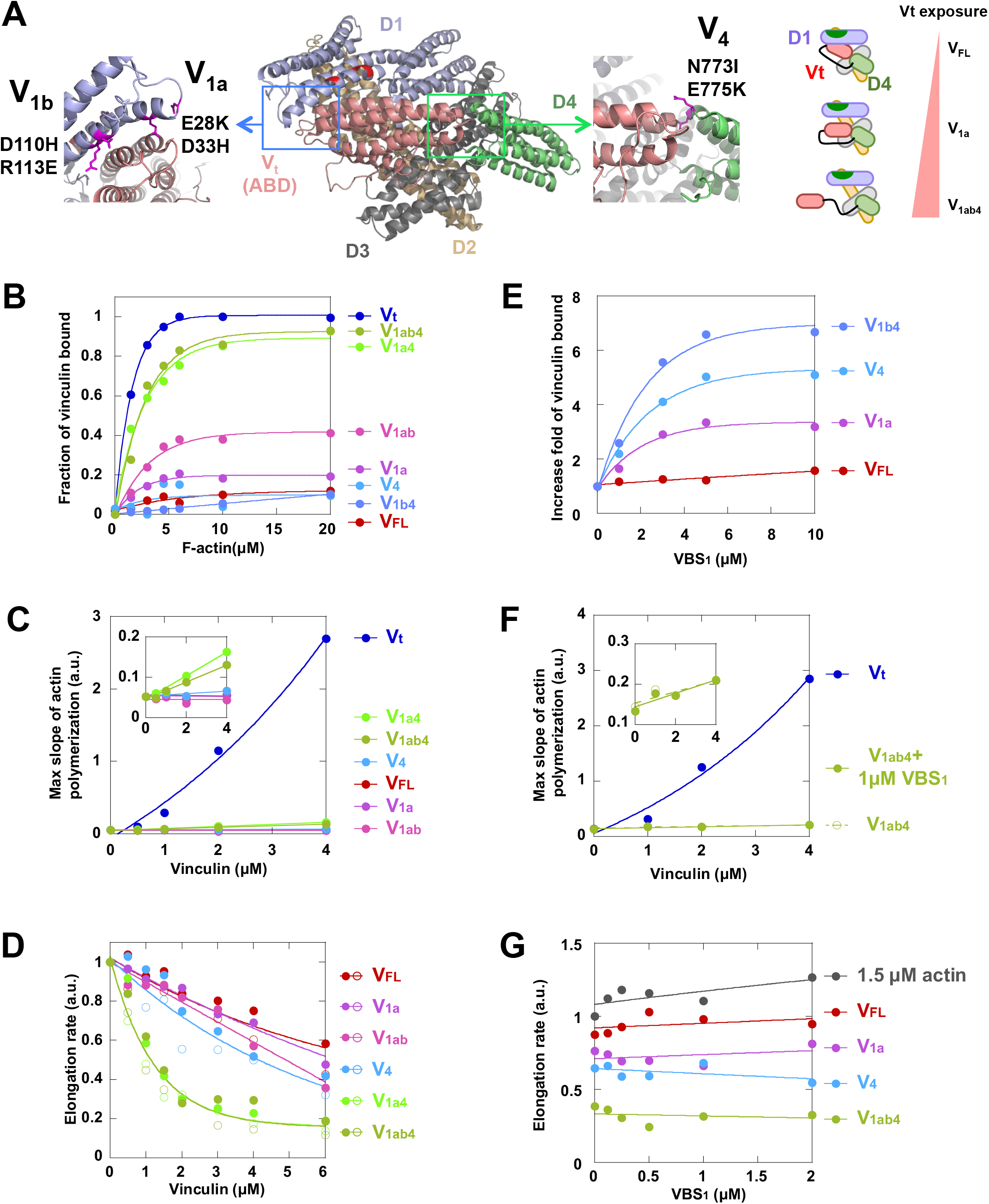
The release of the autoinhibitory contacts of vinculin allows F-actin binding and barbed-end capping but not nucleation. **(A)** Structure of vinculin featuring the subdomains and the auto-inhibitory contacts D1-V_t_ (blue box) and D4-V_t_ (green box). In D1, the double mutations E28K/D33H and D110H/R113E are referred to as V_1a_ and V_1b_ respectively. The double mutation N773I/E775K in D4 is referred to as V_4_. The mutated amino acids are in magenta. A schematic representation of the vinculin mutants is shown on the right. **(B)** Quantification of the cosedimentation of vinculin mutants (2 μM) in the presence of increasing concentration of F-actin. The fraction of vinculin depletion in the supernatants is plotted against F-actin concentration (Supplementary Figure 1). **(C)** The maximal rate of spontaneous actin polymerization (1.5 μM, 10% pyrenyl-labeled) is plotted against the concentration of vinculin mutants (Supplementary Figure 2). V_t_ is used as a positive control for nucleation. The inset is a zoom without V_t_. **(D)** Actin filament barbed end elongation is measured in the presence of spectrin-actin seeds (100 pM), 2 μM actin (10% pyrenyl-labeled) and increasing concentrations of vinculin mutants (Supplementary Figure 3). The fraction of barbed end elongation is the ratio between the elongation rates in the presence and absence of vinculin. Open and closed circle indicate independent experiments in the same conditions. **(E)** The binding of the vinculin mutants (2 μM) to F-actin (10 μM) is measured in a cosedimentation assay in the presence of increasing concentrations of talin VBS_1_ (Supplementary Figure 5). The ratio between vinculin bound to F-actin in the presence and absence of VBS_1_ is plotted against the concentration of VBS_1_. **(F)** The maximal rate of spontaneous actin polymerization (1.5 μM, 10% pyrenyl-labeled) is plotted against the concentration of V_1ab4_ in the absence or presence of 1 μM VBS_1_ (Supplementary Figure 6). V_t_ alone is used as a positive control for nucleation. The inset is a zoom without V_t_. **(G)** Actin filament barbed end elongation is measured in the presence of spectrin-actin seeds (100 pM), 1.5 μM actin (10% pyrenyl-labeled), vinculin mutants (2 μM) and increasing concentrations of VBS_1_ (Supplementary Figure 7). The fraction of barbed end elongation is the ratio of elongation rates in the presence of vinculin and VBS_1_ and in the presence of actin alone.

In order to disrupt specifically both autoinhibitory interfaces, we designed a series of point mutations of amino acids in D1 and D4 that interact with V_t_. Because the contact surface between V_t_ and D1 is significantly larger than the D4-V_t_ one, we mutated two separate groups of charged amino acids in the D1-V_t_ interface: E28K, D33H referred to as V_1a_ and D110H, R113E referred to as V_1b_ (Figure 1A). Their combination is referred to as V_1ab_. In D4, we designed a double mutation N773I, E775K called V_4_ (Figure 1A). In most of our comparative studies, we used full-length vinculin (V_FL_), the isolated ABD V_t_ and the following combinations of point mutants: V_1a_, V_1ab_, V_4_, V_1a4_, V_1b4_, V_1ab4_.

We first characterized these vinculin mutants by measuring their binding to actin filaments (F-actin) using a cosedimentation assay. As previously reported, the autoinhibited V_FL_ does not cosediment with F-actin (Le Clainche et al., 2010) (Figure 1B, Supplementary Figure 1A). Among the mutants, V_1a4_ and V_1ab4_ display the highest affinity for F-actin, close to that of V_t_, which indicates that V_t_ is almost completely exposed in these mutants (Figure 1B, Supplementary Figure 1B, C, D). In contrast V_4_ and V_1b4_ do not bind to F-actin (Figure 1B, Supplementary Figure 1E, F). Finally, V_1a_ and V_1ab_ display low to intermediate binding levels (Figure 1B, Supplementary Figure 1G, H). Altogether, these data demonstrate that the complete release of the D1-V_t_ contact is sufficient to partially expose the F-actin binding site of V_t_, whereas releasing D4 alone is not sufficient. The combined release of D4 and D1 is required to fully expose V_t_ (Supplementary Table 1).

We then tested the effect of these vinculin mutants on actin nucleation by measuring the acceleration of the kinetics of pyrenyl-labeled actin assembly in low ionic strength condition, which corresponds to 25 mM KCl, while the salt concentration normally used by the vast majority of laboratories is 50 mM KCl and sometimes 100 mM KCl. Several studies have revealed regulation mechanisms and activities of FA proteins, such as vinculin, talin and VASP, by varying the ionic strength of in vitro assays (Dedden et al., 2019; Laurent et al., 1999; Le Clainche et al., 2010). We first confirmed that V_t_ nucleates actin filament in a dose-dependent manner (Le Clainche et al., 2010) (Figure 1C, Supplementary Figure 2A). Unlike V_t_, most vinculin mutants do not show significant actin nucleation activities (Figure 1C, Supplementary Figure 2B-G, Supplementary Table 1). Although their activities are very weak, the kinetics showed that V_1a4_ and V_1ab4_ are the only mutants to display a measurable activity (inset in Figure 1C, Supplementary Figure 2F, G), suggesting that the release of V_t_ from both D1 and D4 is necessary to nucleate actin, but not sufficient to fully expose this activity of V_t_.

These two series of experiments lead to an apparent contradiction, since the release of the 2 contacts D1-V_t_ and D4-V_t_ is sufficient to expose V_t_ and allow it to bind to Factin (Figure 1B), but not to nucleate actin filaments (Figure 1C). We hypothesized that, in addition to releasing the D1-V_t_ and D4-V_t_ contacts, F-actin, which is present in cosedimentation assays but absent at the beginning of the nucleation assays, contributes to the opening of vinculin by binding to V_t_. To test this hypothesis, we compared the ability of vinculin mutants to cap preexisting actin filament barbed ends. The effect of vinculin mutants on barbed end capping was first assessed by measuring the inhibition of pyrenyl-labeled actin polymerization in the presence of spectrin-actin seeds (Le Clainche et al., 2010). Although V_FL_, V_1a_, V_1ab_ and V_4_ display only weak activity (Figure 1D, Supplementary Figure 3A-D), V_1a4_ and V_1ab4_ strongly inhibit the elongation of preexisting actin filament barbed ends, in agreement with our hypothesis (Figure 1D, Supplementary Figure 3E, F, Supplementary Table 1).

### An isolated vinculin-binding domain of talin promotes vinculin binding to Factin but has no effect on barbed-end capping and nucleation

Before determining the activity of the talin-vinculin complex, it was necessary to determine whether and how the simple binding of talin to vinculin, independently of the activities of talin ABDs, influences the activities of vinculin described above (Figure 1). We therefore used a short domain of talin, called VBS_1_ (talin 482-636) corresponding to R1 deleted from its last helix, which exposes one VBS (Le Clainche et al., 2010) (Supplementary Figure 4).

We tested the effect of increasing concentrations of VBS_1_ (0-10 μM) on the ability of vinculin mutants to bind to F-actin, using a cosedimentation assay (Figure 1E). VBS_1_ induces the binding of vinculin to F-actin with various efficiencies, depending on mutations in vinculin autoinhibitory contacts. VBS_1_ induces a near two-fold increase in V_FL_ binding to F-actin (Figure 1E, Supplementary Figure 5A). The weakening of D1-V_t_ increases the effect of VBS_1_, which results in a three-fold increase in the binding of V_1a_ to F-actin (Figure 1E, Supplementary Figure 5B). Finally, mutations in D4, alone or combined with mutations in D1, conferred to VBS_1_ its highest efficiency, as shown by the 5-6-fold increase in V_4_ and V_1b4_ binding of F-actin (Figure 1E, Supplementary Figure 5C, D). Taken together, our data show that VBS_1_ binding to D1 mimics mutations in D1, explaining why the combination of VBS_1_ with mutations in D4 mimics the effect of mutations in both D1 and D4 observed previously in Figure 1B (Supplementary Table 1).

VBS_1_ did not increase the nucleation activity of vinculin mutants (Figure 1F, Supplementary Figure 6A-H, Supplementary Table 1), which remains negligible compared to that of V_t_ (Figure 1F, Supplementary Figure 6I). This assay, in which Factin is initially absent, further supports the importance of F-actin as a co-activator of vinculin.

Surprisingly, increasing concentrations of VBS_1_ do not change the ability of vinculin mutants to cap actin filament barbed ends in a spectrin-actin seed assay (Figure 1G, Supplementary Figure 7A-E, Supplementary Table 1). The fact that VBS_1_ does not increase the ability of V_4_ to inhibit barbed end elongation at the same level as V_1ab4_ alone indicates that VBS_1_ binding to D1 is less efficient than D1 mutations in this polymerization assay. The fact that VBS_1_ activates V_4_ in the cosedimentation assay described above (Figure 1E), and not in a kinetic assay (Figure 1G), could be due to the higher concentration of the co-activator F-actin in the cosedimentation assay.

### Talin with exposed VBSs promotes vinculin-dependent barbed-end capping but has no effect on nucleation

After determining the contribution of an isolated talin VBS to the activity of vinculin, we designed a series of full-length talin constructs (T_Δ1_, T_Δ2_, T_Δ3_) in which VBSs are exposed (Figure 2A, Supplementary Figure 4), in order to study the activity of a constitutive talin-vinculin complex harbouring the ABDs of both proteins. We first designed a construct, referred to as T_Δ1_, in which the helix 5 of the helical bundle R1 is deleted, which exposes the same VBS than that of the construct called VBS_1_ used in Figure 1. Talin T_Δ2_ has been designed to expose two consecutive VBSs in R3, thanks to the deletion of the 2 helices 10 and 13 located on both sides of the two VBSs. Talin T_Δ3_ combines the mutations of T_Δ1_ and T_Δ2_. We first verified the efficiency of deletions performed in T_Δ1_, T_Δ2_, T_Δ3_ using a micropatterning-based binding assay adapted form our previous work (Ciobanasu et al., 2015, 2014; Vigouroux et al., 2020). Hence, T_Δ1_, T_Δ2_, T_Δ3_ constructs immobilized on micropatterned surfaces recruit the head of vinculin fused to EGFP (V_h_-EGFP) more efficiently than the wild-type T_FL_ (Figure 2B, C). Because it is impossible to distinguish the relative contribution of each protein in the talin-vinculin complex to F-actin binding, we did not perform cosedimentation assays for this part of the study.

**Figure 2.**
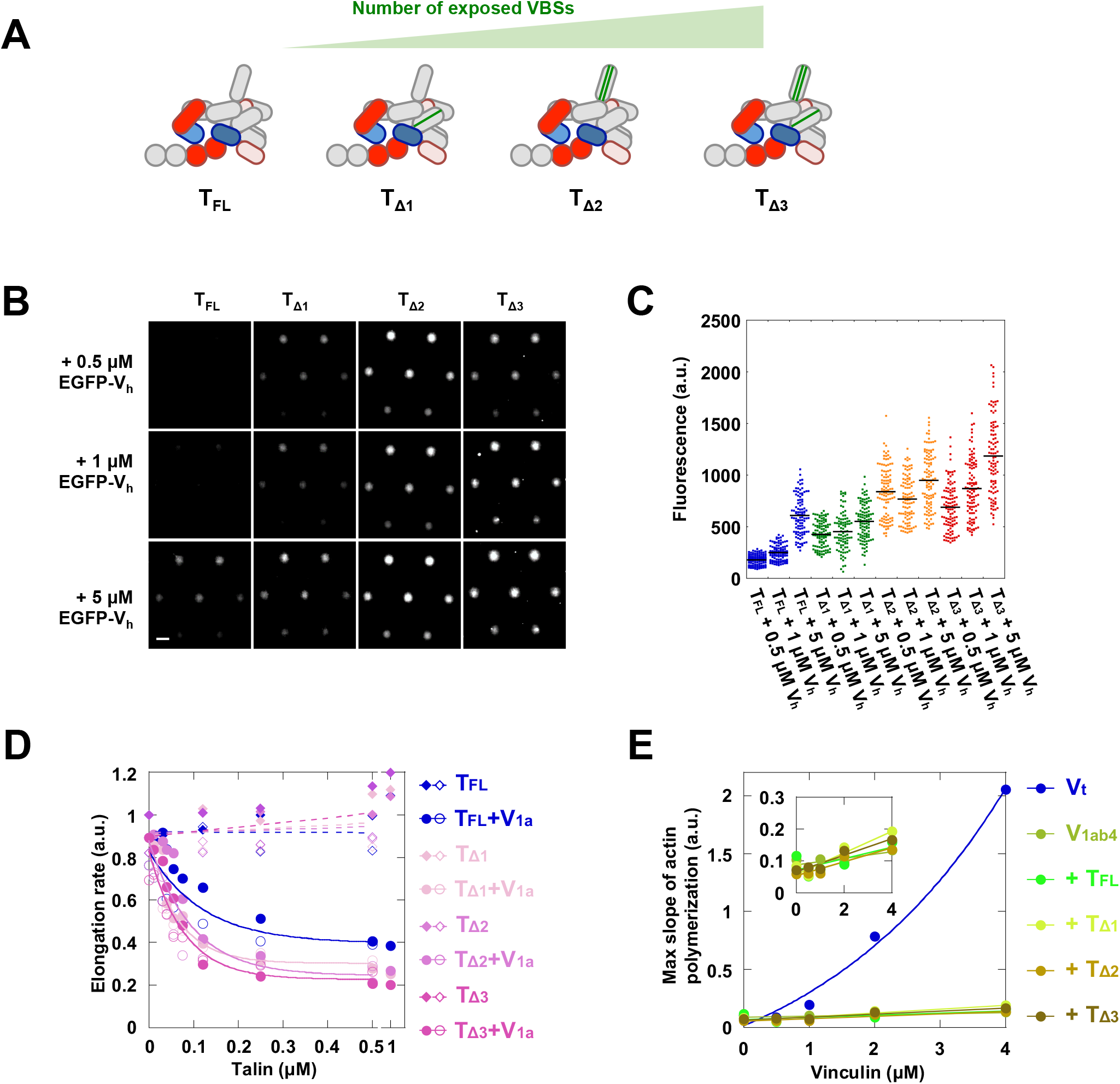
Full-length talin with exposed VBSs promotes vinculin-dependent barbed-end capping but not nucleation. **(A)** Schematic representation of talin constructs with increasing numbers of exposed VBS used in (B-E). **(B, C)** Exposure of VBSs in talin increases the binding of vinculin. **(B)** A micropatterned surface is incubated with 1 μM of the indicated talin contructs, washed, incubated with 0.5 μM, 1 μM and 5 μM EGFP-V_h_ and imaged in TIRF microscopy. Scale bar = 10 μm. **(C)** Quantification of the experiment presented in (A). Each data point represents the mean fluorescence of EGFP-V_h_ in one disk. The bar shows the mean. **(D)** Actin filament barbed end elongation is measured in the presence of spectrin-actin seeds (100 pM), actin (1.5 μM, 10% pyrenyl-labeled) and increasing concentration of talin mutants, in the absence and presence of vinculin V_1a_ (2 μM) (Supplementary Figure 8). The fraction of barbed end elongation was calculated as the ratio of the elongation rate in the presence of increasing concentrations of talin, in the presence or absence of V_1a_, to the elongation rate of actin alone. Note the break between 0.5 μM and 1 μM on the x-axis. Open and closed symbols indicate independent experiments in the same conditions. **(E)** The maximal rate of spontaneous actin polymerization (1.5 μM, 10% pyrenyl-labeled) is plotted against increasing concentrations of V_1ab4_ alone and in the presence of 1 μM of the indicated talin mutants (Supplementary Figure 11). V_t_ alone is used as a positive control for nucleation. The inset is a zoom without V_t_.

We already showed that VBS_1_, which does not include the ABDs of talin, is not sufficient to promote vinculin capping activity (Figure 1G). We then determined the contribution of the ABD domains of talin to the barbed-end capping activity of the talin-vinculin complex. To this aim, we combined the talin mutants with the V_1a_ mutant of vinculin which does not efficiently cap actin barbed ends alone (Figure 1D), nor in the presence of VBS_1_ (Figure 1G). In the presence of a fixed concentration of V_1a_, addition of increasing concentrations of T_Δ1_, T_Δ2_ and T_Δ3_ induced a dose-dependent inhibition of barbed-end elongation in a spectrin-actin seed assay (Figure 2D, Supplementary Figure 8B-D, Supplementary Table 1). Wild-type full-length talin (T_FL_) also induced barbed-end inhibition in the presence of V_1a_, but with a lower efficiency than T_Δ1_, T_Δ2_ and T_Δ3_ (Figure 2D, Supplementary Figure 8A, Supplementary Table 1). This effect is not due to talin constructs alone and requires the presence of V_1a_ since T_FL_, T_Δ1_, T_Δ2_ and T_Δ3_ alone had no effect on barbed end elongation (Figure 2D, Supplementary Figure 9A-D). The fact that the fully autoinhibited V_FL_ does not combine with T_FL_, T_Δ1_, T_Δ2_ and T_Δ3_ to inhibit barbed end elongation confirms that vinculin activation is required for this activity (Supplementary Figure 10A-E).

We then tested the ability of the talin-vinculin complex to stimulate actin assembly by combining the talin mutants with the vinculin mutant V_1ab4_. The mutant V_1ab4_ appeared to be the ideal choice here because it is close to the fully open conformation (Figure 1A). We hypothesized that the weak nucleation activity of V_1ab4_, (Figure 1C, Supplementary Figure 2G), could be increased by talin binding to vinculin head and the activity of talin ABDs. However, kinetic assays containing V_1ab4_ and T_FL_, T_Δ1_, T_Δ2_ or T_Δ3_, do not reveal a significant nucleation activity compared to the activity of V_t_ alone, and it was not higher than that of V_1ab4_ alone (Figure 2E, Supplementary Figure 11A-D, Supplementary Table 1).

### A talin-vinculin complex, in which all autoinhibitory contacts are released, nucleates actin filaments transiently capped at their barbed ends

The results obtained with the previous talin mutants may only partially reflect the activity of the talin-vinculin complex because talin remains autoinhibited by the F3-R9 and F2-R12 contacts (Dedden et al., 2019; Goult et al., 2009). We previously demonstrated that the F3-R9 contact masks the capping activity of talin ABD1 (Ciobanasu et al., 2018). Therefore, we hypothesized that releasing these autoinhibitory contacts in talin may unravel additional activities of the talin-vinculin complex. To test this hypothesis, we designed three talin constructs, called T_ΔAI_, T_Δ1ΔAI_ and T_Δ1ΔAIΔABD2_ (Figure 3A, Supplementary Figure 4). T_ΔAI_ corresponds to T_FL_ deleted from the region encompassing the auto-inhibitory (AI) modules R9 and R12. T_Δ1ΔAI_ combines the deletion of the AI region (ΔAI) and the deletion that exposes a VBS in R1 (Δ1). T_Δ1ΔAIΔABD2_ combines the ΔAI and Δ1 deletions with the deletion of the second actin binding domain (ΔABD2).

**Figure 3.**
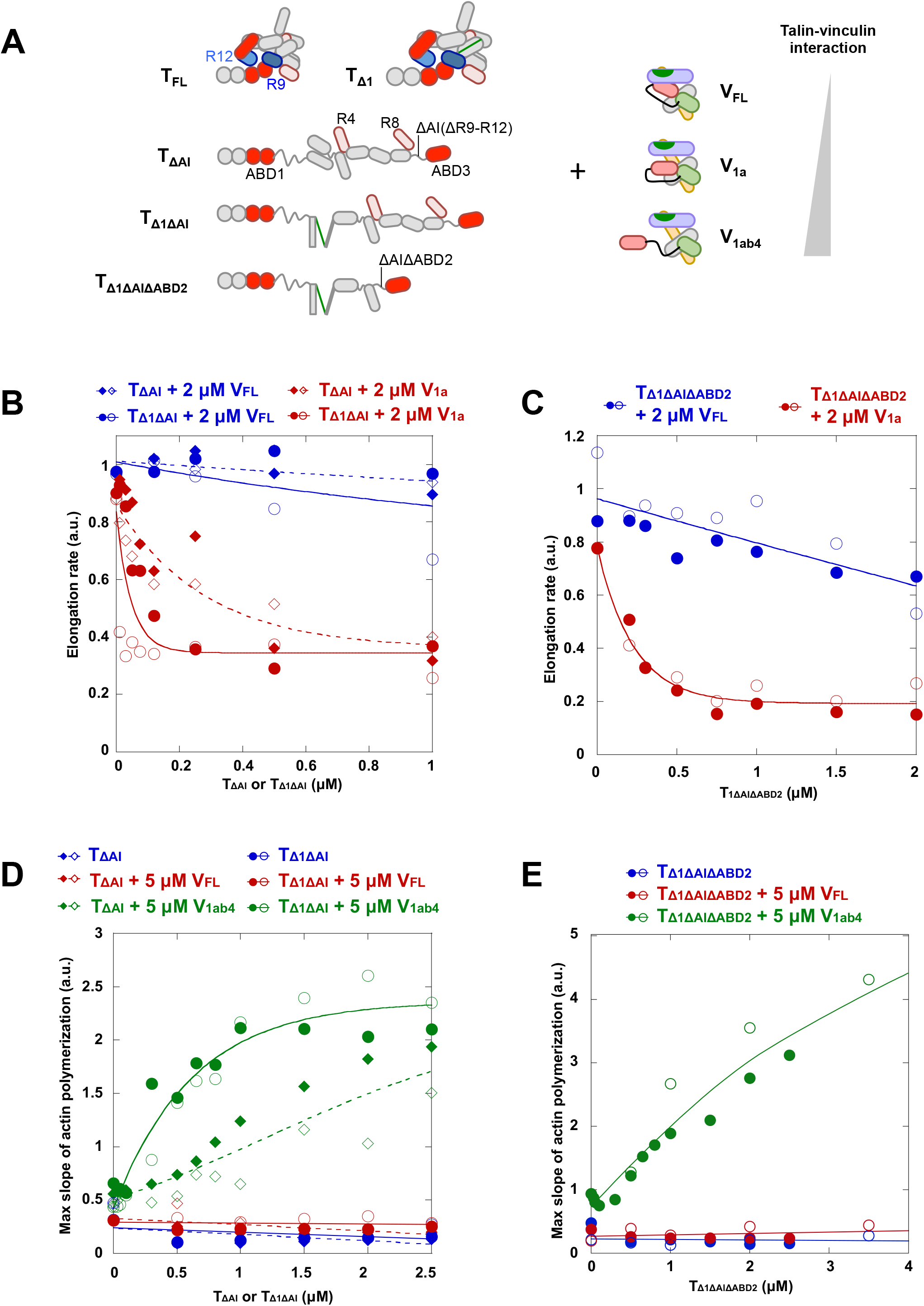
A complex of talin and vinculin, in which all autoinhibitory contacts are released and VBSs are exposed, nucleates and caps actin filaments. **(A)** Schematic representation of the talin and vinculin constructs used in (B-E). **(B, C)** Actin filament barbed end elongation is measured in the presence of 100 pM spectrin-actin seeds, 1.5 μM actin (10% pyrenyl-labeled), 2μM V_FL_ (blue) or 2 μM V_1a_ (red) and increasing concentrations of **(B)** T_ΔAI_ (dash line) or T_Δ1ΔAI_ (solid line) (Supplementary Figure 12), or **(C)** T_Δ1ΔAIΔABD2_ (Supplementary Figure 13). The fraction of barbed end elongation was then calculated as the ratio between the elongation rate in the presence of the indicated combinations of vinculin and talin and the elongation rate of actin alone. **(D, E)** The maximal rate of spontaneous actin polymerization (2.5 μM, 10% pyrenyl-labeled) is plotted as a function of increasing concentrations of **(D)** T_ΔAI_ or T_Δ1ΔAI_, **(E)** T_Δ1ΔAIΔABD2_ alone and in the presence of V_FL_ or V_1ab4_ (Supplementary Figure 15 and 17). (B-E) Open and closed symbols indicate independent experiments in the same conditions.

In a spectrin-actin seed assay, increasing concentrations of T_ΔAI_ and T_Δ1ΔAI_ stimulated the barbed-end capping activity of V_1a_ but not that of V_FL_ (Figure 3B, Supplementary Figure 12A-E, Supplementary Table 1). Altogether these experiments show that releasing the autoinhibitions of talin favors its association with vinculin to form a complex that caps actin filament barbed ends, provided that vinculin autoinhibition is also weakened. The fact that T_Δ1ΔAI_ is much more effective than T_ΔAI_ for this activity further indicates that the release of the autoinhibition of talin adds to the VBS exposure in R1 to promote vinculin binding. The strong barbed-end capping activity of the complex made of T_Δ1ΔAIΔABD2_ and V_1a_, but not V_FL_, also indicates that talin ABD2 is not required for this activity (Figure 3C, Supplementary Figure 13A-C, Supplementary Table 1). The number and position of exposed VBSs influence the barbed end capping activity of the complex since T_Δ2ΔAI_, in which two VBSs are exposed in R3, and T_Δ3ΔAI_, in which three VBSs are exposed in R1 and R3, combine efficiently with V_FL_ to cap actin filaments (Supplementary Figure 14A-E, Supplementary Table 1), in contrast with T_Δ1ΔAI_, in which only one VBS is exposed in R1 (Figure 3B).

We then tested the effect of complexes composed of T_ΔAI_, T_Δ1ΔAI_ and T_Δ1ΔAIΔABD2_ and vinculin mutants on actin nucleation. Increasing concentrations of T_ΔAI_ and T_Δ1ΔAI_ stimulated actin assembly in the presence of V_1ab4_ but not in the presence of V_FL_ (Figure 3D, Supplementary Figure 15A-D, Supplementary Table 1). As for the capping activity (Figure 3B), T_Δ1ΔAI_ is much more effective than T_ΔAI_ for the nucleation activity, indicating that, in addition to the release of talin autoinhibition, VBS exposure is required for talin and vinculin to form a nucleation complex (Figure 3D, Supplementary Figure 15C,D). The number and position of exposed VBSs influence this activity of the complex since a complex made of V_1ab4_ and T_Δ3ΔAI_, in which three VBSs are exposed in R1 and R3, is slightly more active than complexes made of V_1ab4_ and T_Δ1ΔAI_ or T_Δ2ΔAI_, in which one or two VBSs are exposed respectively (Supplementary Figure 16A-F, Supplementary Table 1).

The fact that combinations of activated vinculin V_1ab4_ with auto-inhibited talin (T_Δ1_, T_Δ2_ and T_Δ3_) do not stimulate actin assembly indicates that autoinhibited ABDs in talin play a critical role in this mechanism. To determine the relative importance of the ABDs of talin and vinculin for the activity of the complex, we tested various constructs of the two proteins deleted for ABDs and we added competitors of ABDs to polymerization reactions. First, we showed that talin ABD2 is not necessary since T_Δ1ΔAIΔABD2_, lacking ABD2, efficiently combines with V_1ab4_ to stimulate actin assembly (Figure 3E, Supplementary Figure 17A-C, Supplementary Table 1). We also compared the ability of talin constructs containing VBSs and either ABD1 or ABD3 to stimulate actin assembly in the presence of V_1ab4_. We found that ABD1 and ABD3 can combine with vinculin to stimulate actin polymerization, suggesting redundancy between ABD1 and ABD3 (Supplementary Figure 18A). Interestingly, the addition of the isolated R9 domain of talin to a polymerization reaction containing T_Δ1ΔAIΔABD2_ and V_1ab4_ reduces the stimulation of actin assembly (Supplementary Figure 18B). This result confirms that ABD1, which is masked by R9 in inactive talin (Ciobanasu et al., 2018), is involved in this activity. Finally, combinations of various activated talin constructs with vinculin head (V_h_), which lacks its ABD V_t_, failed to block the elongation of actin filament barbed ends or stimulate actin assembly, indicating a critical role of V_t_ in all activities of the talin-vinculin complex (Supplementary Figure 19A-B). Altogether our observations indicate that ABD1 and ABD3 domains of talin, as well as the V_t_ domain of vinculin, are involved in the activities of the complex.

To confirm the mechanism by which the talin-vinculin complex regulates actin assembly, we observed in TIRF microscopy the actin filaments produced by the complex made of V_1ab4_ and T_Δ1ΔAI_ or T_Δ1ΔAIΔABD2_ described above. Representative images and quantifications showed that the T_Δ1ΔAI_-V_1ab4_ complex induces the formation of a higher number of actin filaments than T_Δ1ΔAI_ alone or V_1ab4_ alone (Figure 4A, C, Supplementary Movie 1). We found the same result for T_Δ1_ΔAIΔABD2-V_1ab4_ complex (Figure 4B, D, Supplementary Movie 2). Interestingly, the analysis of actin filament elongation showed that they undergo frequent pauses reflecting barbed end capping events (Figure 4E, F). The exponential fit of the distribution of the pause duration gave an estimated dissociation rate constant of 0.006 s^-1^ (t_1/2_ = 110 s) for T_Δ1ΔAI_-V_1ab4_ and 0.009 s^-1^ (t_1/2_ = 75 s) for T_Δ1ΔAIΔABD2_-V_1ab4_ (Figure 4G).

**Figure 4.**
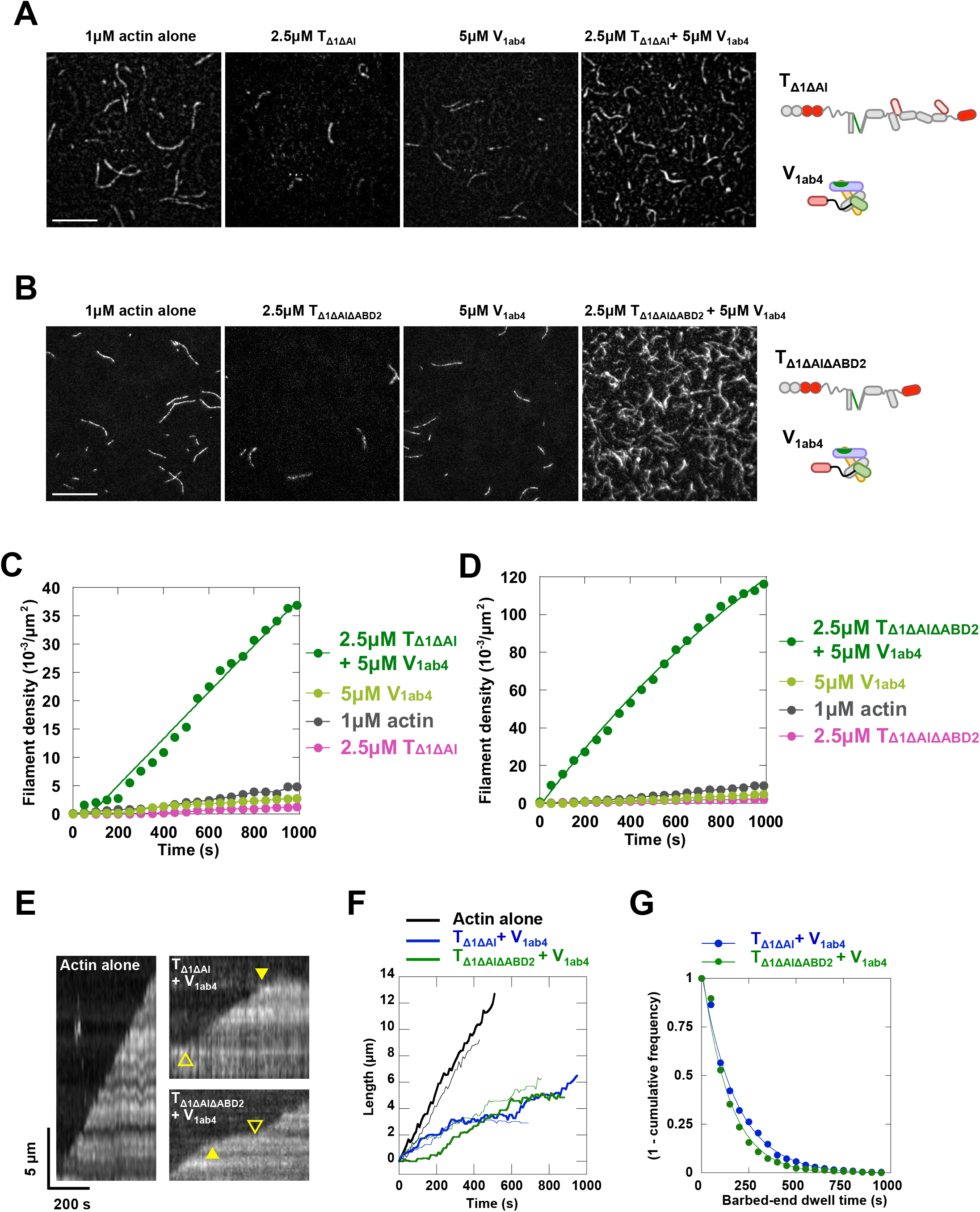
Direct observation of the nucleation and capping of single actin filaments by the talin-vinculin complex. **(A,B)** Single actin filaments observed in TIRF microscopy in the presence of 1 μM actin (5% Alexa488-labeled) alone and supplemented with 2.5 μM T_Δ1ΔAI_ alone, 5 μM V_1ab4_ alone and 2.5 μM T_Δ1ΔAI_ with 5 μM V_1ab4_ **(A),** and in the presence of 1 μM actin (5% Alexa488-labeled) alone and supplemented with 2.5 μM T_Δ1ΔAIΔABD2_ alone, 5 μM V_1ab4_ alone and 2.5 μM T_Δ1ΔAIΔABD2_ with 5 μM V_1ab4_ **(B).** Scale bar = 15 μm. See Supplementary Movie 1 (A) and Supplementary Movie 2 (B). **(C, D)** Quantification of filament density as a function of time from the experiments in (A) and (B). **(E)** Kymographs of single actin filaments in the presence of the indicated proteins from the experiment in (A) and (B). **(F)** Elongation traces of two filaments for each of the conditions indicated from the experiment in (A) and (B). **(G)** Exponential fit of 1-cumulative frequency of barbed end pause times measured in the presence of 1 μM actin, 2.5 μM T _Δ1ΔAI_ with 5 μM V_1ab4_ (blue line, t_1/2_ = 110.35 ± 4.62 s, n = 1023) and 2.5 μM T _Δ1ΔAIΔABD2_ with 5 μM V_1ab4_ (green line, t_1/2_ = 75.37 ± 1.83 s, n = 1619) from experiments in (A) and (B).

Finally, we followed fluorescent molecules of vinculin, talin and actin in real time during the nucleation and capping of individual actin filaments. We first immobilized pre-formed complexes of T_Δ1ΔAI_-Alexa594 and V_1ab4_-Alexa647 at low concentration on a glass surface passivated with PLL-PEG and added actin-Alexa488 under conditions that allowed nucleation and capping. This experimental setup allowed us to capture sequences of events in which spots, containing both talin and vinculin, associate with actin and release a growing actin filament (Figure 5A, Supplementary Movie 3). In this experiment, as in FAs that form in cells, talin, and under certain conditions vinculin, are dimers that assemble with high stoichiometry. The observed spots therefore contain several molecules of T_Δ1ΔAI_ and V_1ab4_ that assemble via the exposed VBS of T_Δ1ΔAI_ but also via other VBSs with lower affinity as indicated by the activity of the T_ΔAI_-V_1ab4_ experiments (Figure 3D). The stationary fiduciary marks along actin filaments indicate that the talin-vinculin complex remains associated with the pointed end while the barbed end is free to grow on the other side (black arrow head, Figure 5A). Interestingly, nucleation events are often characterized by the existence of a delay between actin recruitment by a talin-vinculin complex and the elongation of a filament (Figure 5B, Supplementary Movie 4). In this series of experiments, pre-existing filaments are also captured and capped at their barbed end (Figure 5C, Supplementary Movie 5). Barbed-end capping is transient as many of the capped filaments are eventually left free to elongate after a delay (Figure 5D, Supplementary Movie 6). These activities lead to a distribution of talin-vinculin complexes at the pointed end of the filaments, following their nucleation, at the barbed end of the filaments, following their capping, or on the side of the filaments (Figure 5E). Quantification of the delay between actin recruitment and elongation allows to determine a rate (0.0047 s^-1^, t_1/2_ = 147 s, Figure 5E), that is close to the dissociation rate of the barbed ends from the talin-vinculin complex observed previously (Figure 4G), suggesting that the talin-vinculin complex nucleates filaments initially capped at their barbed end.

**Figure 5.**
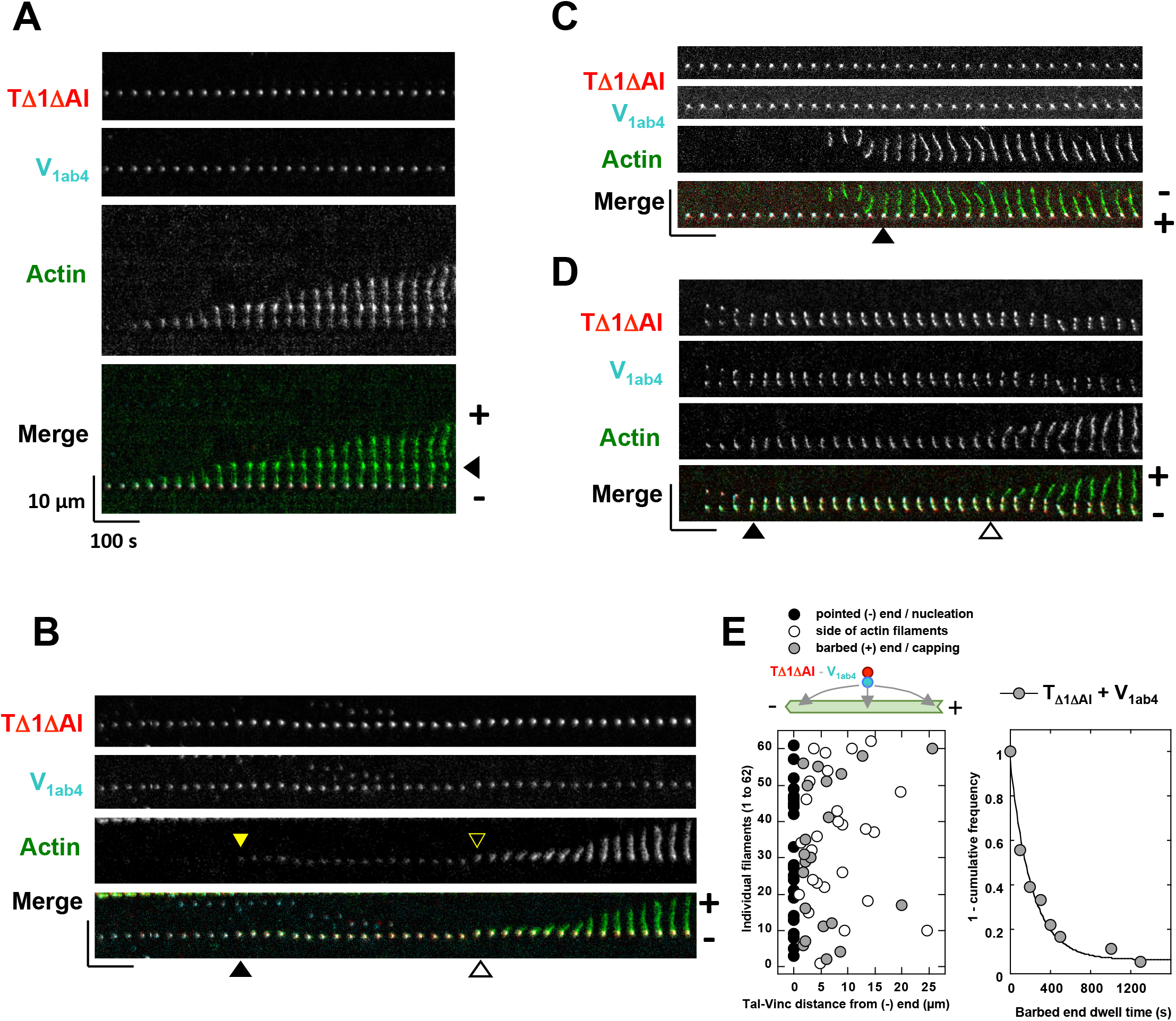
Direct observation of talin, vinculin and actin during the nucleation and barbed-end capping of single actin filaments. **(A-E)** 10 nM T_Δ1ΔAI_ (78% Alexa594-labelled) and 50 nM V_1ab4_ (18% Alexa647-labelled) are immobilized on a glass surface passivated with PLL-PEG, followed by the addition of 0.8 μM actin (5% Alexa488-labelled). **(A)** Kymographs showing the nucleation of an actin filament by a talin-vinculin complex. The stationary fiduciary marks on the right side indicate that the talin-vinculin complex remains associated with the pointed end (-) while the barbed end (+) grows on the other side (Supplementary Movie 3). **(B)** Recruitment of actin by a talin-vinculin complex followed by the elongation of a filament after a delay indicated by the arrowheads (Supplementary Movie 4). The delays are quantified in (E). **(C)** Kymographs showing the capture and barbed-end capping of an existing filament by a talin-vinculin complex (Supplementary Movie 5). **(D)** Kymographs showing the transient barbed-end capping of a filament by a talin-vinculin complex followed by its release (Supplementary Movie 6). **(E)** (Left) Position of talin-vinculin complexes measured along 62 single filaments aligned on their pointed end (-). This quantification reveals that talin-vinculin complexes are found at the pointed end of newly nucleated filaments (black dots, n = 22), along the side of actin filaments (white dots, n = 26) or at the barbed end of capped filaments (grey dots, n = 21). Some of these events are on the same filaments. (Right) Exponential fit of 1-cumulative frequency of the time between actin recruitment by talin-vinculin complexes and barbed end elongation measured in the conditions described above and illustrated in (B), t_1/2_ = 146.69 ± 18.95 s, n = 18.

Altogether, our direct observations reveal a mechanism in which talin and vinculin, in their active conformations, form a complex that nucleates actin filaments transiently capped at their barbed ends before being released and free to elongate (Figure 6). The talin-vinculin complex may also follow part of this mechanism to cap pre-existing filaments (Figure 6).

**Figure 6.**
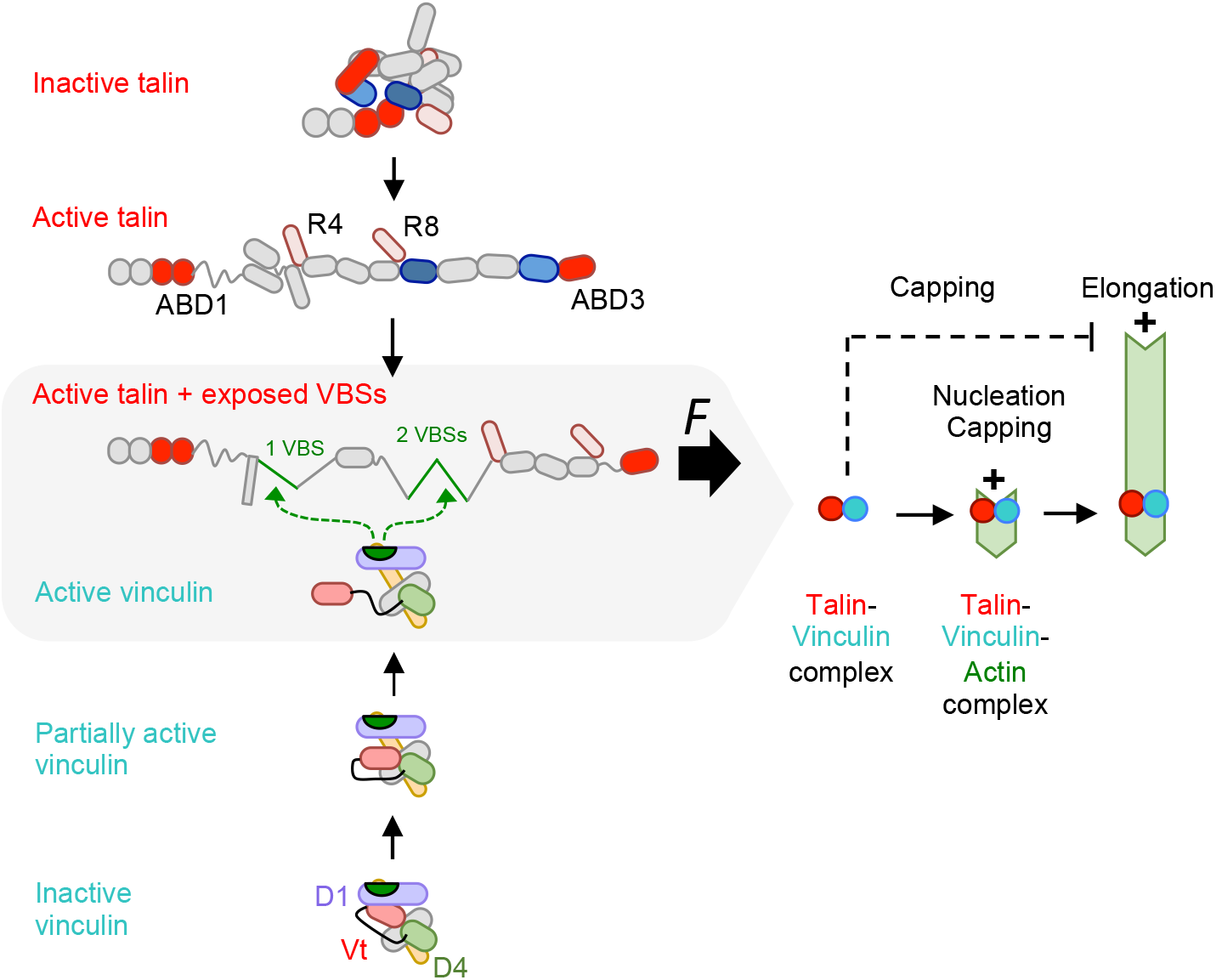
Working model for the activities and regulation of talin, vinculin and the talin-vinculin complex. On the left side, talin (top) and vinculin (bottom) are shown according to their increasing degree of activation towards the central grey box in which the fully activated talin and vinculin form a complex. On the right side, a lower resolution representation shows the activity of the talin-vinculin complex nucleating an actin filament that is transiently capped at its barbed end before being released to elongate.

## Discussion

The assembly of the talin-vinculin complex is a key step in the reinforcement of cellmatrix adhesions subjected to a mechanical stimulus such as the force generated by the actomyosin stress fibers. This complex strengthens the link with the tensile actomyosin cytoskeleton to resist the force it applies. However, the mechanisms by which the force-dependent complex acts on actin filaments was poorly understood.

The mechanistic study of the talin-vinculin complex has long been limited by the fact that the assembly of the complex depends on the release of multiple autoinhibitory interactions both in talin and in vinculin. Although it is established that force exposes cryptic vinculin binding sites in talin by stretching helical bundles, the mechanisms allowing the release of the additional autoinhibitory interactions between distant domains in talin are not fully understood. The structure of full-length talin in its autoinhibited form has been reported recently and revealed that, in addition to the known interaction between F3 and R9, the F2 domain interacts with the R12 domain (Dedden et al., 2019; Goult et al., 2009). These autoinhibitory contacts mask the binding interface of ABD1, corresponding to F2 and F3, with F-actin, while ABD2, composed of R4 and R8, is masked by an interaction with R3 (Atherton et al., 2020; Ciobanasu et al., 2018; Dedden et al., 2019) (Figure 6). The mechanism by which these contacts are disrupted is unclear. The binding of RIAM and PIP2 to talin F3 is known to disrupt the F3-R9 autoinhibition (Yang et al., 2014). Once these two interdomain interactions are disrupted, talin can be pulled by actomyosin force, which stretches helical bundles to expose cryptic VBSs (del Rio et al., 2009; Yao et al., 2016) (Figure 6). Force application to the ABD3/R13 domain of talin could also contribute to disrupting its auto-inhibitions, but no experimental evidence of such a mechanism has been reported to date. This mechanism is possible because the ABD3 domain is exposed, at least partially, in the inactive talin, as suggested by the flexibility of the neighboring domains (Dedden et al., 2019). In this study, we show that combining various deletions, including the part of talin that contains the R9 and R12 autoinhibitory domains, with the deletion of individual helices, which exposes VBSs, mimics the open and mechanically stretched conformation of talin (Figure 6).

Our results show that the release of talin autoinhibition is sufficient to allow the binding of vinculin and form a complex that caps and nucleates actin, which explains the observation made by others that a mutation in talin that disrupts the F3-R9 interaction is sufficient to recruit vinculin in cells (Atherton et al., 2020). Our data also support the findings that the association of talin and vinculin without tension is required for efficient nascent adhesion maturation (Han et al., 2021). This interaction likely corresponds to a low degree of saturation of the partially exposed VBSs in this open talin structure that has become very flexible. Indeed, exposure of a single VBS motif in R1 significantly increases vinculin binding as evidenced by the enhanced activity of the talin-vinculin complex.

Vinculin is autoinhibited by an intramolecular interaction between the head (V_h_) and the tail (V_t_) domains. Talin binding to the D1 subdomain of V_h_ is controlled by the D1-V_t_ interaction (Johnson and Craig, 1994), whereas F-actin binding to V_t_ is controlled by a more complex mechanism involving interactions of D1 and D4 with both sides of V_t_ (Cohen et al., 2005; Johnson and Craig, 1995; Le Clainche et al., 2010). The identification of several intermediate open conformations of vinculin suggests that the level of activation of vinculin is controlled by the number of auto-inhibitory contacts removed (Chorev et al., 2018). Our vinculin mutagenesis strategy confirmed that V_t_ activities are controlled by different combinations of V_h_-V_t_ autoinhibitory contacts. The disruption of D1-V_t_ is sufficient to induce F-actin binding to V_t_, but the combination of D1 and D4 mutations provides a higher affinity for F-actin. In the present study we show that the disruption of D1-V_t_ is not sufficient to expose the capping activity of V_t_, which requires the combined disruption of D1 and D4 contacts with V_t_, as previously suggested (Le Clainche et al., 2010). Early studies showed that the coincidental binding of talin and F-actin to D1 and V_t_ respectively keep vinculin in its open conformation (Chen et al., 2006). In agreement with this model, full-length vinculin dissociates rapidly from FAs after actomyosin inhibition, whereas V_h_ remains stably associated, demonstrating that tensile actomyosin stress fibers bound to V_t_ maintain the open conformation of talin-bound vinculin (Carisey et al., 2013; Humphries et al., 2007). This observation is further explained by the discovery that V_t_ forms a catch bond with F-actin in response to force (Huang et al., 2017). It is therefore not surprising that the release of D1 and D4 from V_t_ in our experiments releases the capping activity of vinculin that occurs in the presence of actin filaments, but results in very little actin nucleation initiated without actin filaments.

Solving the problem of releasing the auto-inhibitory contacts of talin and vinculin was a prerequisite for determining the activity that results from the combination of their ABDs. It is interesting to remember that the combination of actin-binding proteins within transient complexes is often found in important signal-responsive mechanisms for regulating actin dynamics. However, the results of such combinations cannot be predicted from the activities of the isolated proteins, as exemplified by the formation of branched actin filaments resulting from the combination of WASP and Arp2/3 (Le Clainche and Carlier, 2008). Here, we show that the combination of talin ABDs with vinculin V_t_ in a complex creates a machinery that nucleates actin filaments transiently capped at their barbed end (Figure 6). In this mechanism, talin and vinculin cooperate to form a new filament through a mechanism that is not yet understood but which probably involves the stabilization of actin nuclei that form spontaneously in solution by the ABDs of talin and vinculin. This mechanism leads to the transient barbed-end capping of the newly formed actin filaments. After being released from capping, the actin filament elongates by its free barbed end, while the complex remains attached to the side of the filament, near the pointed end (Figure 6). It would be tempting to attribute to vinculin alone the nucleation and capping activities of the talin-vinculin complex, since V_t_ has these activities (Le Clainche et al., 2010; Wen et al., 2009). However, our previous report indicates that the capping activity of the complex involves the ABD1 of talin (Ciobanasu et al., 2018). The fact that a complex, composed of active vinculin and talin with exposed VBSs but masked ABD1, does not nucleate, shows unambiguously the importance of ABD1 for nucleation. Although the function of the ABD1 domain of talin is generally restricted to integrin activation in a membrane-bound conformation that does not allow its binding to actin, it is important to recall here that talin is in equilibrium between actin and the plasma membrane as evidenced by its retrograde movement in FAs (Hu et al., 2007). ABD2 is not involved in capping or nucleation mechanisms as demonstrated by the strong activities of a mutant that does not contain this domain. However, ABD3 can also combine with V_t_ to nucleate, suggesting a possible redundancy of ABD1 and ABD3 in this mechanism. It is interesting to see that the three domains of talin are redundant for the bundling of actin filaments (Kelley et al., 2020), but are involved more selectively in the actin polymerization activities of talin in combination with vinculin. It is very likely that actin filaments nucleated by talin and vinculin will end up forming large bundles, but our microscopy experiments, where the filaments are short and spaced out, disadvantages this activity and we have not explored this direction already widely documented by recent studies (Boujemaa-Paterski et al., 2020; Kelley et al., 2020).

In the context of a stress fiber anchored to a focal adhesion, where this talin-vinculin complex resides, it is unlikely that this release of actin filaments, occurring with a half-time of 75 s to 110 s, allows the fast processive elongation of a stress fiber. However, this release is fast enough to feed stress fibers with new actin filaments and reinforce stress fibers. It would be interesting to determine how the talin-vinculin complex cooperates with other ABPs in FAs, such as VASP, parvin or tensin, to regulate actin assembly.

## Material and methods

### Recombinant cDNA Constructs

cDNAs encoding for vinculin 1-1066 (V_FL_), vinculin E28K/D33H (V_1a_), vinculin E28K/D33H/D110H/R113E (V_1ab_), vinculin N773I/E775K (V_4_), vinculin E28K/D33H/N773I/E775K (V_1a4_), vinculin D110H/R113E/N773I/E775K (V_1b4_), vinculin E28K/D33H/D110H/R113E/N773I/E775K (V_1ab4_) were synthezied and subcloned in the NcoI site of pET-3d by Genscript. cDNAs encoding for vinculin 879-1066 (V_t_), talin-1 482–636 (VBS_1_), talin-1 1655–1822 (R9), full-length talin-1 (T_FL_), full-length vinculin (V_FL_) were described previously (Ciobanasu et al., 2018, 2015; Le Clainche et al., 2010). cDNAs encoding for T_Δ1ΔAIΔABD2_, T_F2F3R1R2R3_, T_R2R3ABD3_ were constructed by different PCR amplifications of the full length talin cDNA and subcloning in a pETM plasmid. N-terminal Strep-tag II, PreScission site and C-terminal His_6_ tag were introduced by PCR. cDNAs of T_ΔAI_, T_Δ1_, T_Δ2_, T_Δ3_, T_Δ1ΔAI_, T_Δ2ΔAI_, T_Δ3ΔAI_ were constructed and subcloned into pET-29a+ by Genscript. The constructs were verified by sequencing. See Supplementary Figure 4 for the detail of the talin constructs.

### Protein Purification

V_FL_, V_t_, V_1a_, V_1ab_, V_4_, V_1a4_, V_1b4_, V_1ab4_, T_FL_, T_ΔAI_, T_Δ1_, T_Δ2_, T_Δ3_, T_Δ1ΔAI_, T_Δ2ΔAI_, T_Δ3ΔAI_, T_Δ1ΔAIΔABD2_, T_F2F3R1R2R3_, T_R2R3ABD3_ and VBS_1_ were expressed in *E.Coli* BL21 as previously described (Ciobanasu et al., 2015). Briefly, 1mM isopropyl β-D-1-thiogalactopyranoside (IPTG) was used for induction. Bacterial pellets of V_FL_, V_t_, V_1a_, V_1ab_, V_4_, V_1a4_, V_1b4_, V_1ab4_ were lysed by sonication in 20 mM Tris, pH 8.0, 1 M NaCl, 1 mM β-mercaptoethanol, 10 μg/ml benzamidine and 1 mM PMSF. Lysates of His-tagged vinculin constructs were purified by Ni-NTA-sepharose affinity chromatography (Ni^2+^-nitrilotriacetic acid, Qiagen), followed by a Q-Sepharose ion exchange column. V_t_ was purified as previously described (Le Clainche et al., 2010). Vinculin proteins were finally dialyzed in 20 mM Tris, pH 7.8, 1mM DTT. Bacterial pellets of T_FL_, T_ΔAI_, T_Δ1_, T_Δ2_, T_Δ3_, T_Δ1ΔAI_, T_Δ2ΔAI_, T_Δ3ΔAI_, T_Δ1ΔAIΔABD2_, T_F2F3R1R2R3_, T_R2R3ABD3_ were lysed by sonication in 50 mM Tris, pH 7.8, 500 mM NaCl, 1% Triton X-100, 1 mM β-mercaptoethanol, 10 μg/ml benzamidine and 1 mM PMSF. Lysates of His-tagged talin constructs were purified by Ni-NTA-sepharose affinity chromatography (Ni^2+^-nitrilotriacetic acid, Qiagen), followed by a gel filtration column (Superdex 200, 16/600, GE Healthcare). VBS_1_ was purified as previously described (Le Clainche et al., 2010). Talin constructs were finally dialyzed in 20 mM Tris, pH 7.8, 100 mM KCl, 1 mM DTT.

### F-actin co-sedimentation assay

F-actin co-sedimentation assays were performed to determine the affinity of vinculin constructs for F-actin in the absence and presence of talin VBS_1_. In F-actin co-sedimentation assay with vinculin alone, 2 μM of vinculin (V_FL_, V_t_, V_1a_, V_1ab_, V_4_, V_1a4_, V_1b4_, V_1ab4_) was incubated with 0, 1.5, 3, 4.5, 6, 10, 20 μM F-actin in 5 mM Tris, pH 7.8, 100 mM KCl, 1 mM MgCl_2_, 0.2 mM EGTA, 200 μM ATP, 1 mM DTT during 15 min at room temperature. To test the effect of VBS_1_, F-actin co-sedimentation assays were also performed in the presence of 2 μM of vinculin (V_FL_, V_1a_, V_4_, V_1b4_), 10 μM Factin, with or without 1, 3, 5, 10 μM VBS_1_ in 5 mM Tris, pH 7.8, 100 mM KCl, 1 mM MgCl_2_, 0.2 mM EGTA, 200 μM ATP, 1 mM DTT during 15 min at room temperature. After centrifugation at 90,000 rpm in a TLA-120.1 rotor (Beckman) during 30 min, the pellets and supernatants were separated and loaded on SDS-PAGE. Gels were scanned and analyzed with the ImageJ software.

### Polymerization assay

Actin polymerization was measured by the increase in fluorescence of 10 % pyrenyl-labeled actin in a SAFAS Xenius spectrofluorimeter (Safas, Monaco). To measure the elongation of actin filament barbed ends, actin polymerization was induced by 100 pM spectrin-actin seeds in 10 % pyrenyl-labeled CaATP-G-actin, 100 mM KCl, 1 mM MgCl_2_, and 0.2 mM EGTA in presence of indicated proteins. The fraction of barbed end elongation was calculated as the ratio between the elongation rates in the presence and absence of proteins of interest. To test the ability of proteins to nucleate actin filaments, spontaneous polymerization was induced by adding 25 mM KCl, 1 mM MgCl_2_, and 0.2 mM EGTA to 10% pyrenyl-labeled CaATP-G-actin solution in presence of the proteins of interest.

### Observation of talin, vinculin and single actin filaments in TIRF microscopy

Our protocol is a modification of protocols used to study talin and vinculin activities (Ciobanasu et al., 2018; Le Clainche et al., 2010). To prevent the nonspecific binding of actin filaments to the surface of the coverslip, we first irradiated coverslips with deep UVs for 3 min and incubated them with 0.1 mg/mL PLL-PEG for 1 h at room temperature. The coverslip was then washed extensively with water and dried. Flow cells containing 40-60 μl of liquid were prepared by sticking the PLL-PEG-coated coverslip to a slide with double-sided adhesive spacers. In experiments to observe and count single filament number, the chamber was first incubated with washing buffer (5 mM Tris, pH 7.8, 200 μM ATP, 1 mM DTT, 1 mM MgCl_2_, 0.2 mM CaCl_2_, 25 mM KCl) for 1 min. The chamber was then saturated with 10% BSA for 5 min and washed with washing buffer. The final reaction was then injected into the chamber. A typical reaction was composed of 1 μM actin (5% Alexa488-labeleld) in 5 mM Tris, pH 7.8, 200 μM ATP, 1% methylcellulose, 5 mM 1,4-diazabicyclo(2,2,2)-octane, 25 mM KCl, 1 mM MgCl_2_, 200 μM EGTA, 40 mM DTT supplemented with various talin and vinculin mutants. To observe talin, vinculin and actin simultaneously, 0.2 μM T_Δ1ΔAI_ (78% Alexa594-labelled) and 1 μM V_1ab4_ (18% Alexa647-labelled) were first mixed, diluted 20 times to reach final concentrations of 10 nM T_Δ1ΔAI_ and 50 nM V_1ab4_, injected on a flow chamber to immobilize T_Δ1ΔAI_-V_1ab4_ complexes non-specifically on the surface passivated with PLL-PEG, and finally supplemented with 0.8 μM actin (5% Alexa488-labelled). Finally, we sealed the flow chamber with VALAP (a mixture of Vaseline, lanolin, and paraffin) and observed the reaction on an Nikon Eclipse Ti-E inverted microscope equipped with 60X/1.49NA objective (Nikon). The time-lapse videos were acquired by Metamorph and subsequently analyzed by the ImageJ software.

### Protein labeling

For 3-color imaging, actin is labeled with Alexa Fluor 488 carboxylic acid, succinimidyl ester, which reacts with the NH_2_ groups of basic amino acids (Ciobanasu et al., 2015). T_Δ1ΔAI_ and V_1ab4_ are labeled on free cysteines with Alexa Fluor 594 maleimide and Alexa Fluor 647 maleimide respectively.

### Quantification and statistical analysis

All experiments presented in this manuscript were repeated independently two to four times. Where appropriate, data points from two independent experiments are overlaid to demonstrate reproducibility. For unambiguous negative results, a single data set is presented.

## Acknowledgements

This work was supported by the Agence Nationale de la Recherche grants ANR-16-CE13-0007-02 PHAGOMECANO and ANR ANR-18-CE13-0026-01 RECAMECA (to CLC). HW is supported by a PhD fellowship from the China Scholarship Council (CSC). RS is supported by a PhD fellowship from Association de Spécialisation et d’Orientation Scientifique. The present work has benefited from the Light Microscopy facility of Imagerie-Gif, (http://www.i2bc.paris-saclay.fr), member of IBiSA (http://www.ibisa.net), supported by “France-BioImaging” (ANR-10-INBS-04-01), and the Labex “Saclay Plant Sciences” (ANR-10-LABX-0040-SPS). We thank the members of the “Cytoskeleton Dynamics and Motility” team for helpful discussions.

## Author contributions

HW performed the binding assays, the kinetic assays and the microscopy experiments, purified most of the proteins, analyzed the data and prepared the figures. RS performed additional kinetic assays. VH designed and cloned some of the cDNAs. CV performed binding assays on micropatterned surfaces. JP contributed to single filament microscopy assays. CLC designed the experiments, supervised the project and wrote the manuscript.

## Legends of supplementary figures and movies

**Supplementary figure 1. The high affinity binding of vinculin to the side of actin filaments requires the disruption of the contacts made by V_t_ with both D1 and D4. (A-H)** SDS-PAGE gels showing the supernatant and pellet fractions of cosedimentation assays containing 2 μM of the indicated vinculin mutants and increasing concentrations of F-actin (0, 1.5, 3, 4.5, 6, 10, 20 μM).

**Supplementary figure 2. The release of the D1-V_t_ and D4-V_t_ contacts does not allow actin nucleation by vinculin. (A-G)** Spontaneous actin polymerization was measured in the presence of increasing concentrations of the indicated vinculin mutants and 1.5 μM actin (10% pyrenyl-labeled) in a low salt buffer (25 mM KCl). The control kinetics corresponding to 1.5 μM actin alone are the same in all panel.

**Supplementary figure 3. The release of the D1-V_t_ and D4-V_t_ contacts allows barbed-end capping by vinculin. (A-F)** The elongation of actin filament barbed end was measured in the presence of increasing concentrations of the indicated vinculin mutants, 100 pM spectrin-actin seeds, 2 μM actin (10% pyrenyl-labeled). The control kinetics showing the polymerization of 2 μM actin alone are the same in all panels.

**Supplementary figure 4. Talin constructs used in the study.** Domain organisation of full-length talin and talin constructs featuring intramolecular autoinhibitions in red, VBSs in green and ABDs in pink. Grey bars indicate α-helices in the rod. Red stars indicate exposed VBSs.

**Supplementary figure 5. Talin VBS_1_ combined with D4-V_t_ release induces vinculin binding to F-actin. (A-D)** SDS-PAGE gels showing the pellet fractions of Factin co-sedimentation reactions containing 2 μM of the indicated vinculin mutants, 10 μM F-actin and increasing concentrations of talin VBS_1_ (0, 1, 3, 5, 10 μM).

**Supplementary figure 6. Talin VBS_1_ combined with D4-V_t_ release has no effect on nucleation by vinculin. (A-I)** Spontaneous actin polymerization was measured in the presence of increasing concentrations of the indicated vinculin mutants and 1.5 μM actin (10% pyrenyl-labeled) in a low salt buffer (25 mM KCl) in the absence of VBS_1_ **(A-D, I)** and in presence of 1 μM VBS_1_ **(E-H)**. The control kinetics showing the polymerization of 1.5 μM actin in the presence of 1 μM V_t_, as a positive control, or in the presence of 1 μM VBS_1_ alone, as a negative control, are the same in all panels.

**Supplementary figure 7. Talin VBS_1_ combined with D4-V_t_ release has no effect on barbed-end capping by vinculin. (A-E)** The elongation of actin filament barbed end was measured in the presence of 2 μM of the indicated vinculin mutants with increasing concentrations of VBS_1_ (0.12, 0.25, 0.5, 1, 2 μM), 100 pM spectrin-actin seeds and 1.5 μM actin (10% pyrenyl-labeled). The control kinetics showing the polymerization of 1.5 μM actin alone are the same in all panels.

**Supplementary figure 8. A complex composed of full-length talin with exposed VBSs and V_1a_ caps actin filament barbed ends. (A-D)** The elongation of actin filament barbed end was measured in the presence of 2 μM V_1a_ with increasing concentrations of the indicated talin mutants, 100 pM spectrin-actin seeds and 1.5 μM actin (10% pyrenyl-labeled). The control kinetics showing the polymerization of 1.5 μM actin in the presence of 2 μM V_1a_ are the same in all panels.

**Supplementary figure 9. Talin mutants with exposed VBSs alone do not cap barbed ends. (A-D)** The elongation of actin filament barbed end was measured in the presence of increasing concentrations of the indicated talin mutants, 100 pM spectrin-actin seeds, 1.5 μM actin (10% pyrenyl-labeled). The kinetics with 1.5 μM G-actin, 2 μM V_FL_ or 2 μM V_1a_, used here as additional negative controls, are the same in all panels.

**Supplementary figure 10. Addition of full-length talin with exposed VBSs to V_FL_ does not cap actin filament barbed ends. (A-D)** The elongation of actin filament barbed end was measured in the presence of 2 μM V_FL_, increasing concentrations of the indicated talin mutants, 100 pM spectrin-actin seeds, 1.5 μM actin (10% pyrenyl-labeled). The kinetics with 1.5 μM actin and 2 μM V_FL_ are the same in all panels. **(E)** The fraction of barbed end elongation was calculated as the ratio of the elongation rate in the presence of V_FL_ and talin mutants to the elongation rate of 1.5 μM actin alone.

**Supplementary figure 11. Talin mutants with exposed VBSs combined with V_1ab4_ do not stimulate actin assembly. (A-D)** Spontaneous actin polymerization was measured in the presence of 1μM of the indicated talin mutants, increasing concentrations of V_1ab4_ and 1.5 μM actin (10% pyrenyl-labeled) in a low salt buffer (25 mM KCl). The kinetics with 1.5 μM actin and 1 μM V_t_ alone, used as a positive control for nucleation, or 1.5 μM actin and 4 μM V_1ab4_ alone are the same in all panels.

**Supplementary figure 12. Talin mutants, with released autoinhibitory contacts and one exposed VBS combine with V_1a_ to cap actin filament barbed ends.** The elongation of actin filament barbed end was measured in the absence and presence of 2 μM V_FL_ or 2 μM V_1a_ or 1 μM T_ΔAI_ or 1 μM T_Δ1ΔAI_ **(A)**, in the presence of increasing concentrations of indicated talin mutants and 2 μM V_FL_ **(B-C)** or 2 μM V_1a_ **(D-E)**, 100 pM spectrin-actin seeds and 1.5 μM actin (10% pyrenyl-labeled). The control kinetics with 1.5 μM actin and 2 μM V_FL_ **(A-C)** or 1.5 μM actin and 2 μM V_1a_ **(D-E)** are the same in all panels.

**Supplementary figure 13. A talin mutant, with released autoinhibitory contacts, one exposed VBS and lacking ABD2, combines efficiently with V_1a_, but not with V_FL_, to cap actin filament barbed ends. (A-C)** The elongation of actin filament barbed end was measured in the presence of increasing concentrations of T _Δ1ΔAIΔABD2_, 100 pM spectrin-actin seeds and 1.5 μM actin (10% pyrenyl-labeled) (A), increasing concentrations of T _Δ1ΔAIΔABD2_, 2 μM V_FL_ (B) or 2 μM V_1a_ (C), 100 pM spectrin-actin seeds and 1.5 μM actin (10% pyrenyl-labeled). The kinetics with 1.5 μM actin alone and 1.5 μM actin and V_t_ are the same in all panels.

**Supplementary figure 14. Talin mutants, with released autoinhibitory contacts and two or three exposed VBSs, combine with V_FL_ and V_1a_ to cap actin filament barbed ends. (A-E)** The elongation of actin filament barbed end was measured in the presence of increasing concentrations of T_Δ2ΔAI_ and 2 μM of V_FL_ **(A)**, increasing concentrations of T_Δ2ΔAI_ and 2 μM of V_1a_ **(B)**, increasing concentrations of T_Δ3ΔAI_ and 2 μM of V_FL_ **(C)**, increasing concentrations of T_Δ3ΔAI_ and 2 μM of V_1a_ **(D)**, 100 pM spectrin-actin seeds and 1.5 μM actin (10% pyrenyl-labeled). The control kinetics with 1.5 μM actin and 2 μM V_t_ or 1 μM T_Δ2ΔAI_ or 1 μM T_Δ3ΔAI_ or 2 μM V_FL_ or 2 μM V_1a_ are the same in all panels. **(E)** Quantification of the experiments in (A-D).

**Supplementary figure 15. Talin mutants, with released autoinhibitory contacts and one exposed VBS, combine efficiently with V_1ab4_ to stimulate actin nucleation. (A-D)** Actin polymerization was measured in the absence and presence of 5 μM of V_FL_ or V_1ab4_ with increasing concentrations of T_ΔAI_ or T_Δ1ΔAI_ and 2.5 μM G-actin (10% pyreyl-labeled) in a low salt buffer (25 mM KCl). The kinetics with 2.5 μM actin alone, used as a negative control, and 2.5 μM actin and 5 μM V_t_, used as a positive control for nucleation, are the same in all panels.

**Supplementary figure 16. Talin mutants, with released autoinhibitory contacts and two or three exposed VBSs, combine efficiently with V_1ab4_ to stimulate actin nucleation. (A-D)** Actin polymerization was measured in the presence of 5 μM of V_FL_ or V_1ab4_ with increasing concentrations of T_Δ2ΔAI_ or T_Δ3ΔAI_ and 2.5 μM actin (10% pyreyl-labeled) in a low salt buffer (25 mM KCl). **(E)** Additional negative control showing the polymerization of 2.5 μM actin (10% pyreyl-labeled) in the presence of 5 μM V_FL_ or 5 μM V_1ab4_ or 1.5 μM T_Δ2ΔAI_ or 2.5 μM T_Δ3ΔAI_. The kinetics with 2.5 μM actin alone, used as a negative control, and 2.5 μM actin and 5 μM V_t_, used as a positive control for nucleation, are the same in all panels. **(F)** The maximal rate of spontaneous actin polymerization is plotted against increasing concentrations of talin mutants in the presence of V_FL_ or V_1ab4_ as described in (A-D). Data for T_Δ1ΔAI_ + V_FL_ or V_1ab4_ and T_ΔAI_ + V_FL_ or V_1ab4_ are the same as in Figure 3D.

**Supplementary figure 17. A talin mutant, with released autoinhibitory contacts, one exposed VBS and lacking ABD2, combines efficiently with V_1ab4_ to stimulate actin nucleation. (A-C)** Spontaneous actin polymerization was measured in the presence of 2.5 μM actin (10% pyreyl-labeled) and increasing concentrations of T_Δ1ΔAIΔABD2_ alone (A), increasing concentrations of T_Δ1ΔAIΔABD2_ and 5 μM V_FL_ (B) and increasing concentrations of T_Δ1ΔAIΔABD2_ and 5 μM V_1ab4_ (C) in a low salt buffer (25 mM KCl). The kinetics with 2.5 μM actin and 5 μM V_t_ alone, used as a positive control or 5 μM V_1ab4_ alone are the same in all panels.

**Supplementary figure 18. Importance of talin ABDs for the activities of the talin-vinculin complex. (A,B)** Spontaneous actin polymerization was measured in presence of 2.5 μM actin (10% pyrenyl-labeled) and the indicated talin and vinculin mutants in a low salt buffer (25 mM KCl). The black arrows in (B) indicate the stimulation of actin polymerization by V_1ab4_ alone and V_1ab4_ + T_Δ1ΔAIΔABD2_. The dark green arrow in (B) indicates that R9 reverses the actin polymerization induced by V_1ab4_ + T_Δ1ΔAIΔABD2_.

**Supplementary figure 19. A talin-vinculin complex, in which vinculin lacks V_t_, does not nucleate and cap actin filaments. (A)** The elongation of actin filament barbed end was measured in the presence of the indicated proteins, 100 pM spectrin-actin seeds, 1.5 μM actin (10% pyrenyl-labeled). **(B)** Spontaneous actin polymerization was measured in presence of 2.5 μM actin (10% pyrenyl-labeled) and the indicated proteins in a low salt buffer (25 mM KCl).

**Supplementary Movie 1. Observation of the activity of T_Δ1ΔAI_ and V_1ab4_ on single actin filaments in TIRF microscopy.** Conditions: 1 μM actin (5% Alexa 488-labeled) in 5 mM Tris, pH 7.8, 200 μM ATP, 1% methylcellulose, 5 mM 1,4-diazabicyclo(2,2,2)-octane (DABCO), 25 mM KCl, 1 mM MgCl_2_, 200 μM EGTA, 40 mM DTT supplemented with 2.5 μM T_Δ1ΔAI_ or 5 μM V_1ab4_ or 2.5 μM T_Δ1ΔAI_ and 5 μM V_1ab4_. Scale bar = 15 μm.

**Supplementary Movie 2. Observation of the activity of T_Δ1ΔAIΔABD2_ and V_1ab4_ on single actin filaments in TIRF microscopy.** Conditions: 1 μM actin (5% Alexa 488-labelled) in 5 mM Tris, pH 7.8, 200 μM ATP, 1% methylcellulose, 5 mM 1,4-diazabicyclo(2,2,2)-octane (DABCO), 25 mM KCl, 1 mM MgCl_2_, 200 μM EGTA, 40 mM DTT supplemented with 2.5 μM T_Δ1ΔAIΔABD2_ or 5 μM V_1ab4_ or 2.5 μM T_Δ1ΔAIΔABD2_ and 5 μM V_1ab4_. Scale bar = 15 μm.

**Supplementary Movie 3. Observation of talin, vinculin and actin during the nucleation of a single actin filament.** 0.2 μM T_Δ1ΔAI_ (78% Alexa594-labelled) and 1 μM V_1ab4_ (18% Alexa647-labelled) were first mixed, diluted 20 times to reach final concentrations of 10 nM T_Δ1ΔAI_ and 50 nM V_1ab4_, injected on a flow chamber to immobilize T_Δ1ΔAI_-V_1ab4_ complexes non-specifically on the surface, and finally supplemented with 0.8 μM actin (5% Alexa488-labelled).

**Supplementary Movie 4. Recruitment of actin by a talin-vinculin complex followed by the elongation of a filament after a delay.** 0.2 μM T_Δ1ΔAI_ (78% Alexa594-labelled) and 1 μM V_1ab4_ (18% Alexa647-labelled) were first mixed, diluted 20 times to reach final concentrations of 10 nM T_Δ1ΔAI_ and 50 nM V_1ab4_, injected on a flow chamber to immobilize T_Δ1ΔAI_-V_1ab4_ complexes non-specifically on the surface, and finally supplemented with 0.8 μM actin (5% Alexa488-labelled).

**Supplementary Movie 5. Capture and barbed-end capping of an existing filament by a talin-vinculin complex.** 0.2 μM T_Δ1ΔAI_ (78% Alexa594-labelled) and 1 μM V_1ab4_ (18% Alexa647-labelled) were first mixed, diluted 20 times to reach final concentrations of 10 nM T_Δ1ΔAI_ and 50 nM V_1ab4_, injected on a flow chamber to immobilize T_Δ1ΔAI_-V_1ab4_ complexes non-specifically on the surface, and finally supplemented with 0.8 μM actin (5% Alexa488-labelled).

**Supplementary Movie 6. Transient barbed-end capping of a filament by a talin-vinculin complex followed by its release.** 0.2 μM T_Δ1ΔAI_ (78% Alexa594-labelled) and 1 μM V_1ab4_ (18% Alexa647-labelled) were first mixed, diluted 20 times to reach final concentrations of 10 nM T_Δ1ΔAI_ and 50 nM V_1ab4_, injected on a flow chamber to immobilize T_Δ1ΔAI_-V_1ab4_ complexes non-specifically on the surface, and finally supplemented with 0.8 μM actin (5% Alexa488-labelled).

**Supplementary Table 1:**
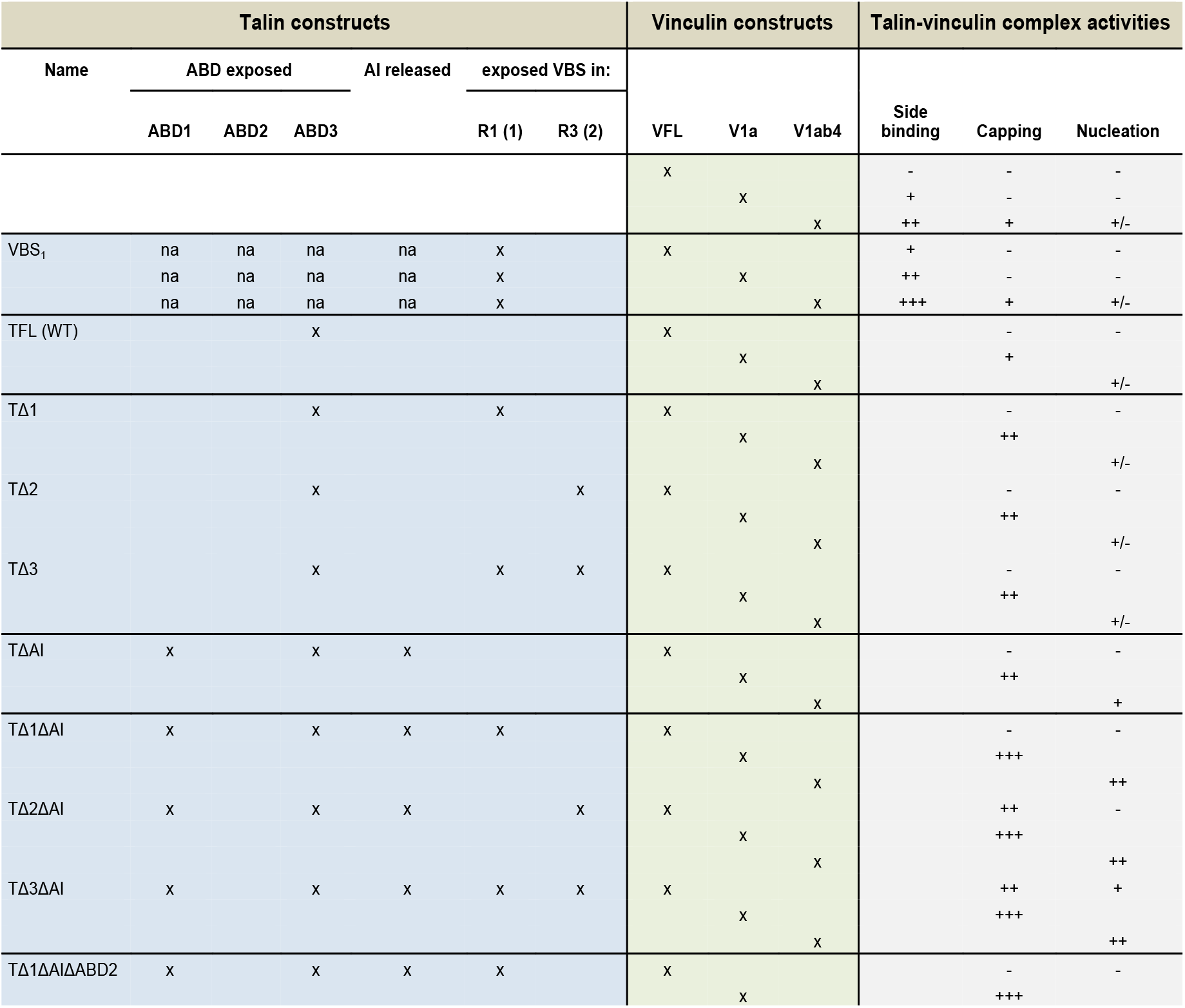
Summary of the activities of talin and vinculin mutants and their complexes measured in kinetic assays.

