## Supplementary Figures 1-19 for "Talin and vinculin combine their activities to trigger actin assembly"

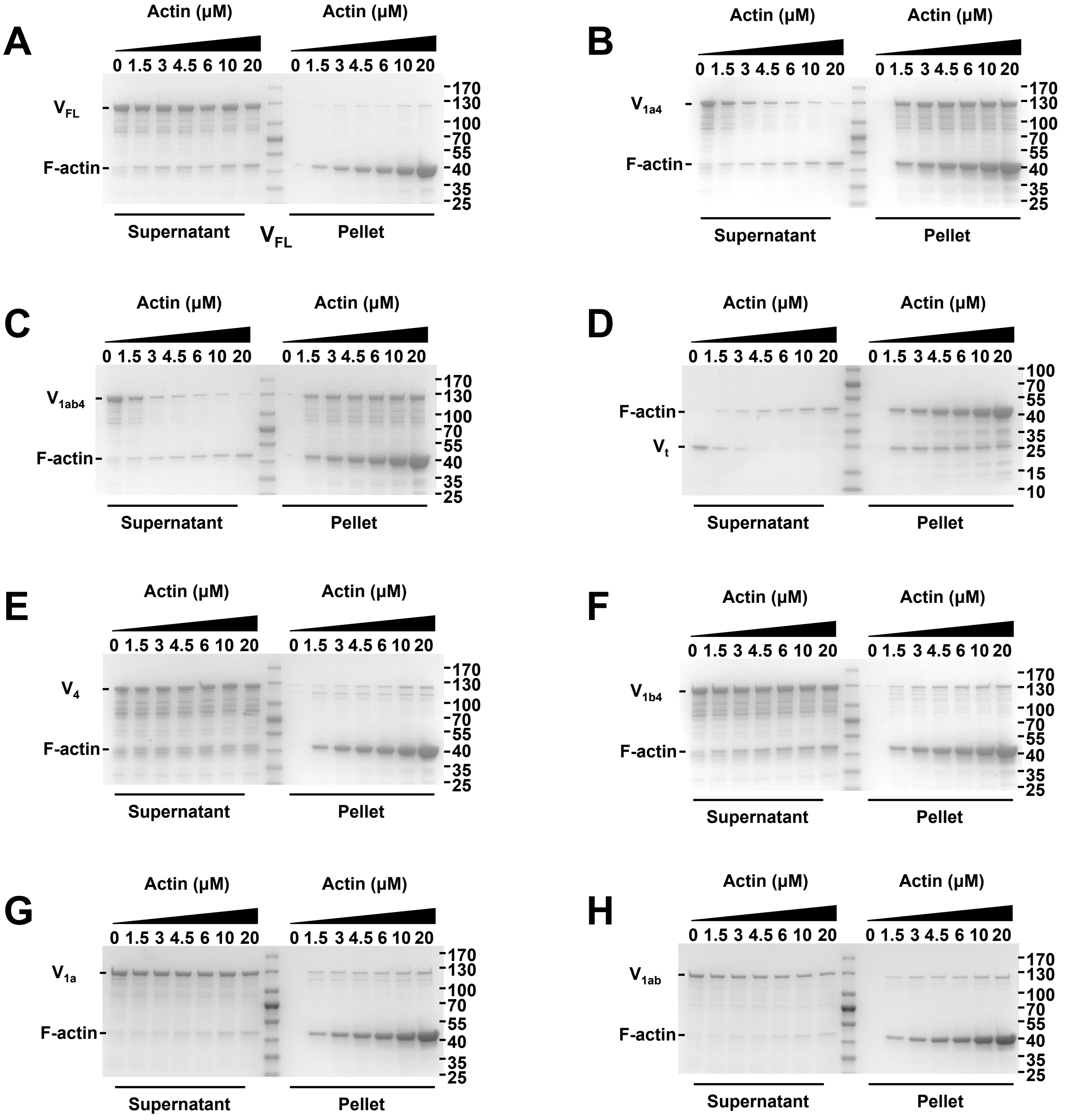


**Supplementary figure 1. The high affinity binding of vinculin to the side of actin filaments requires the disruption of the contacts made by $V_t$ with both D1 and D4. (A-H)** SDS-PAGE gels showing the supernatant and pellet fractions of cosedimentation assays containing 2 μM of the indicated vinculin mutants and increasing concentrations of F-actin (0, 1.5, 3, 4.5, 6, 10, 20 μM).

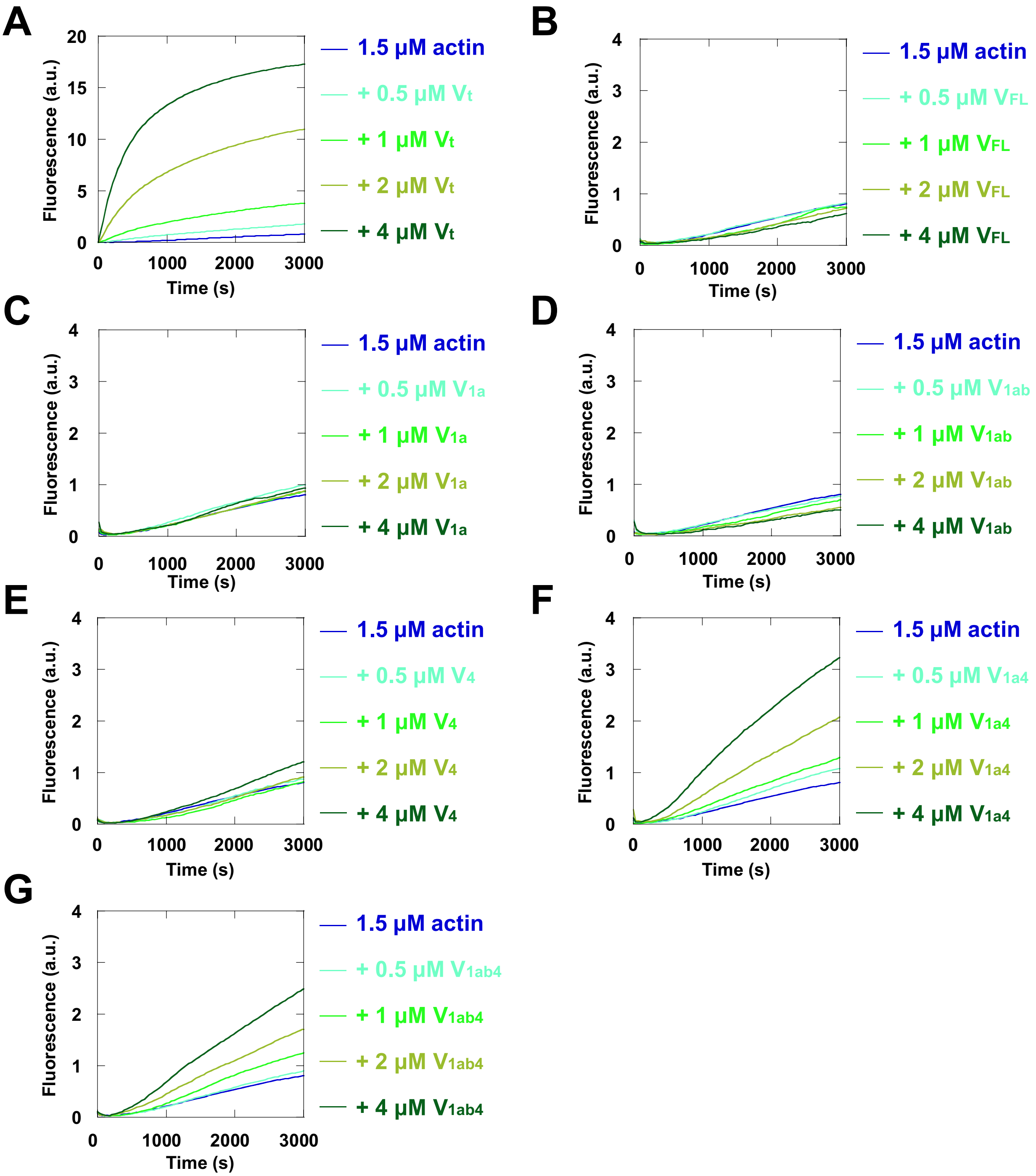


**Supplementary figure 2. The release of the D1-$V_t$ and D4-$V_t$ contacts does not allow actin nucleation by vinculin. (A-G)** Spontaneous actin polymerization was measured in the presence of increasing concentrations of the indicated vinculin mutants and 1.5 μM actin (10% pyrenyl-labeled) in a low salt buffer (25 mM KCl). The control kinetics corresponding to 1.5 μM actin alone are the same in all panel.

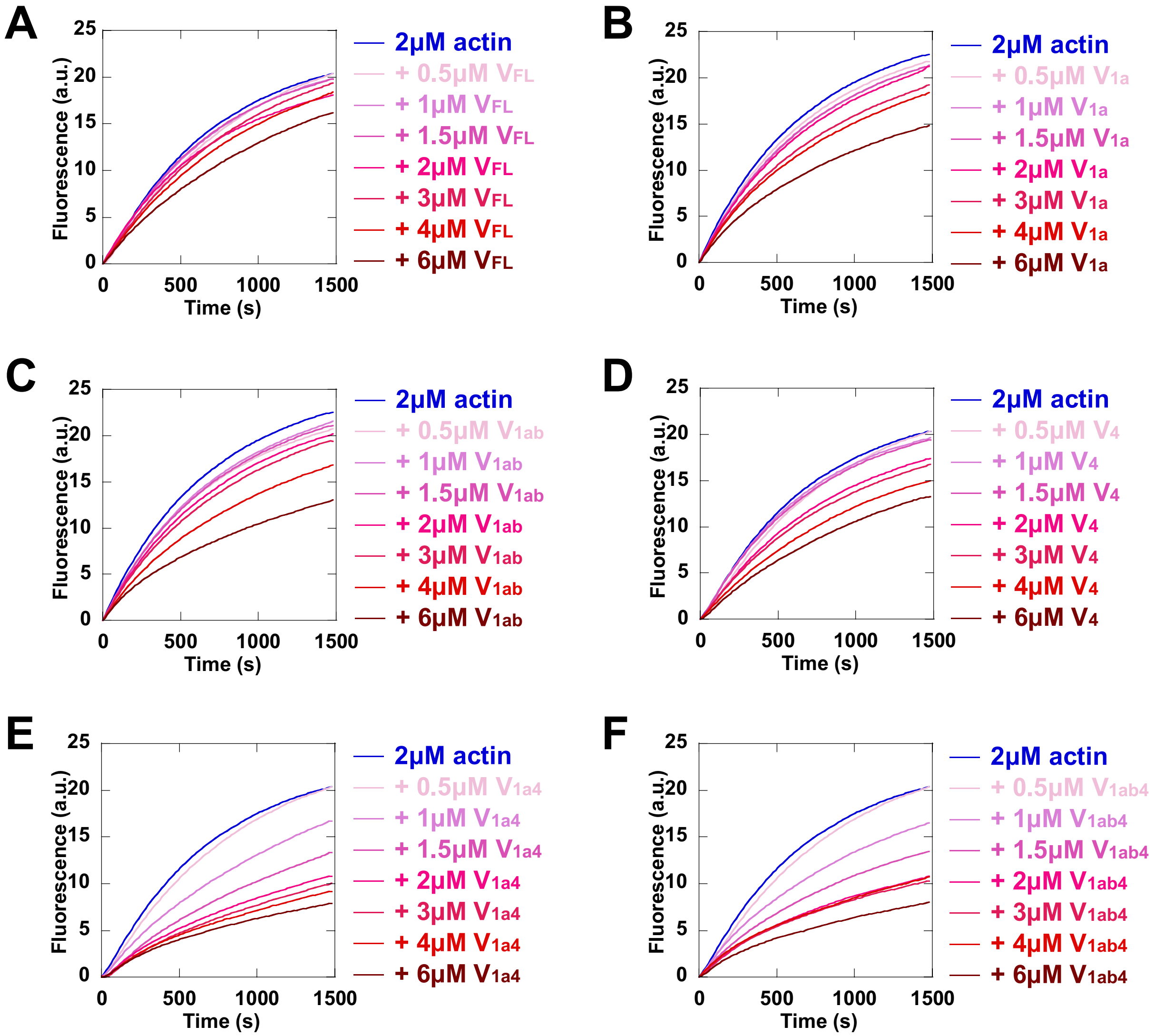


**Supplementary figure 3. The release of the D1-$V_t$ and D4-$V_t$ contacts allows barbed-end capping by vinculin. (A-F)** The elongation of actin filament barbed end was measured in the presence of increasing concentrations of the indicated vinculin mutants, 100 pM spectrin-actin seeds, 2 μM actin (10% pyrenyl-labeled). The control kinetics showing the polymerization of 2 μM actin alone are the same in all panels.

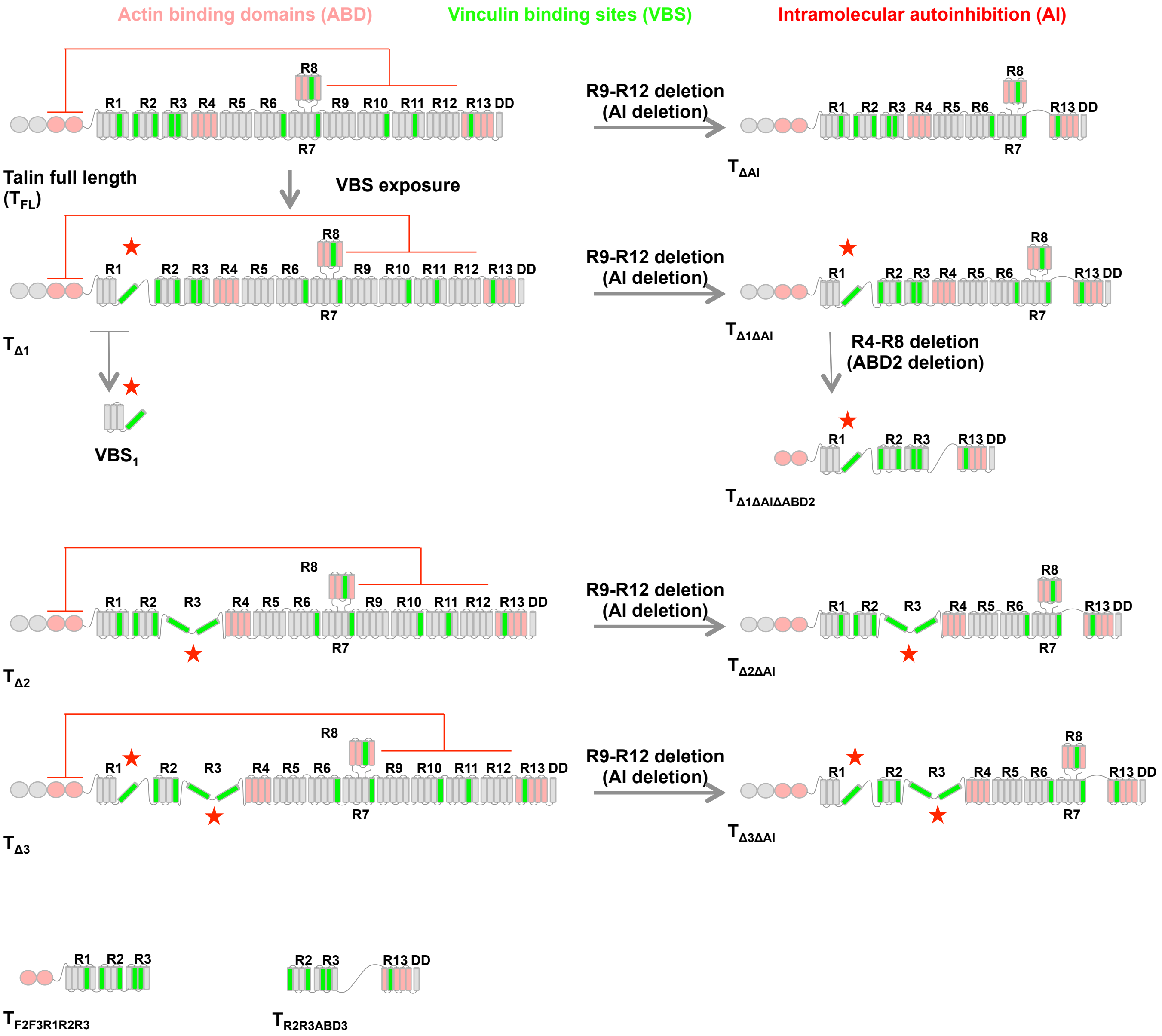


**Supplementary figure 4. Talin constructs used in the study.** Domain organisation of full-length talin and talin constructs featuring intramolecular autoinhibitions in red, VBSs in green and ABDs in pink. Grey bars indicate $\alpha$-helices of the rod. Red stars indicate exposed VBSs.

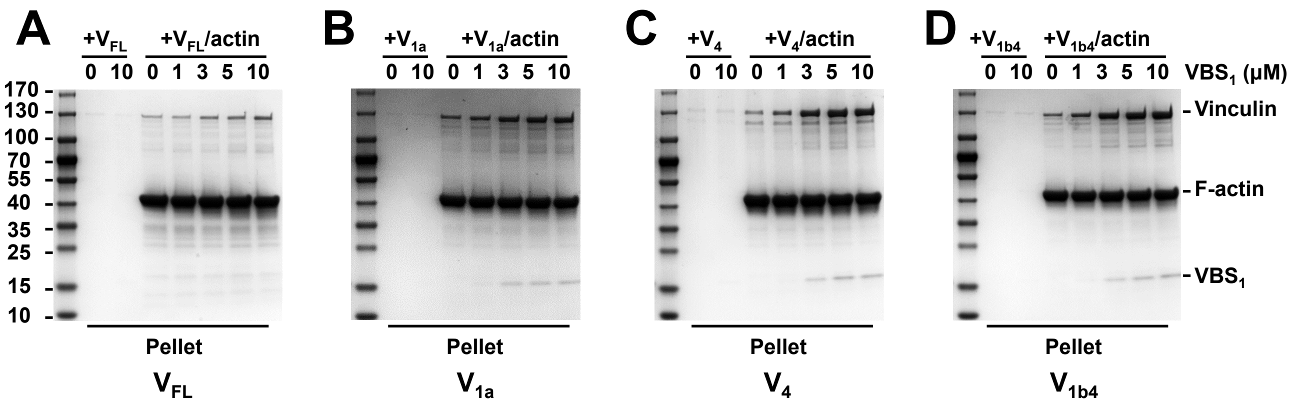


**Supplementary figure 5. Talin $VBS_1$ combined with D4-$V_t$ release induces vinculin binding to F-actin. (A-D)** SDS-PAGE gels showing the pellet fractions of F-actin co-sedimentation reactions containing 2 μM of the indicated vinculin mutants, 10 μM F-actin and increasing concentrations of talin $VBS_1$ (0, 1, 3, 5, 10 μM).

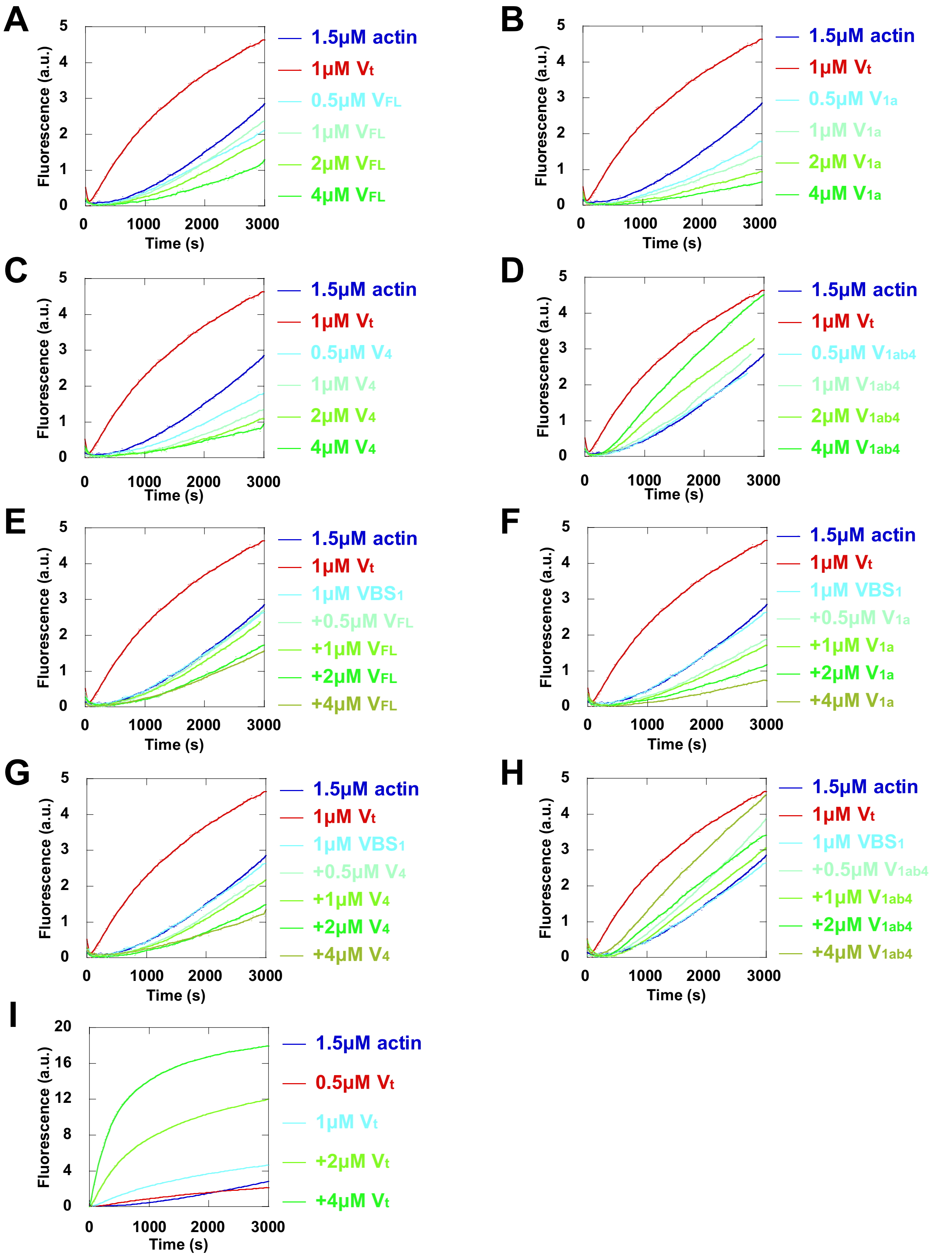


**Supplementary figure 6. Talin VBS$_1$ combined with D4-V$_t$ release has no effect on nucleation by vinculin. (A-I)** Spontaneous actin polymerization was measured in the presence of increasing concentrations of the indicated vinculin mutants and 1.5 μM actin (10% pyrenyl-labeled) in a low salt buffer (25 mM KCl) in the absence of VBS$_1$ **(A-D, I)** and in presence of 1 μM VBS$_1$ **(E-H)**. The control kinetics showing the polymerization of 1.5 μM actin in the presence of 1 μM V$_t$, as a positive control, or in the presence of 1 μM VBS$_1$ alone, as a negative control, are the same in all panels.

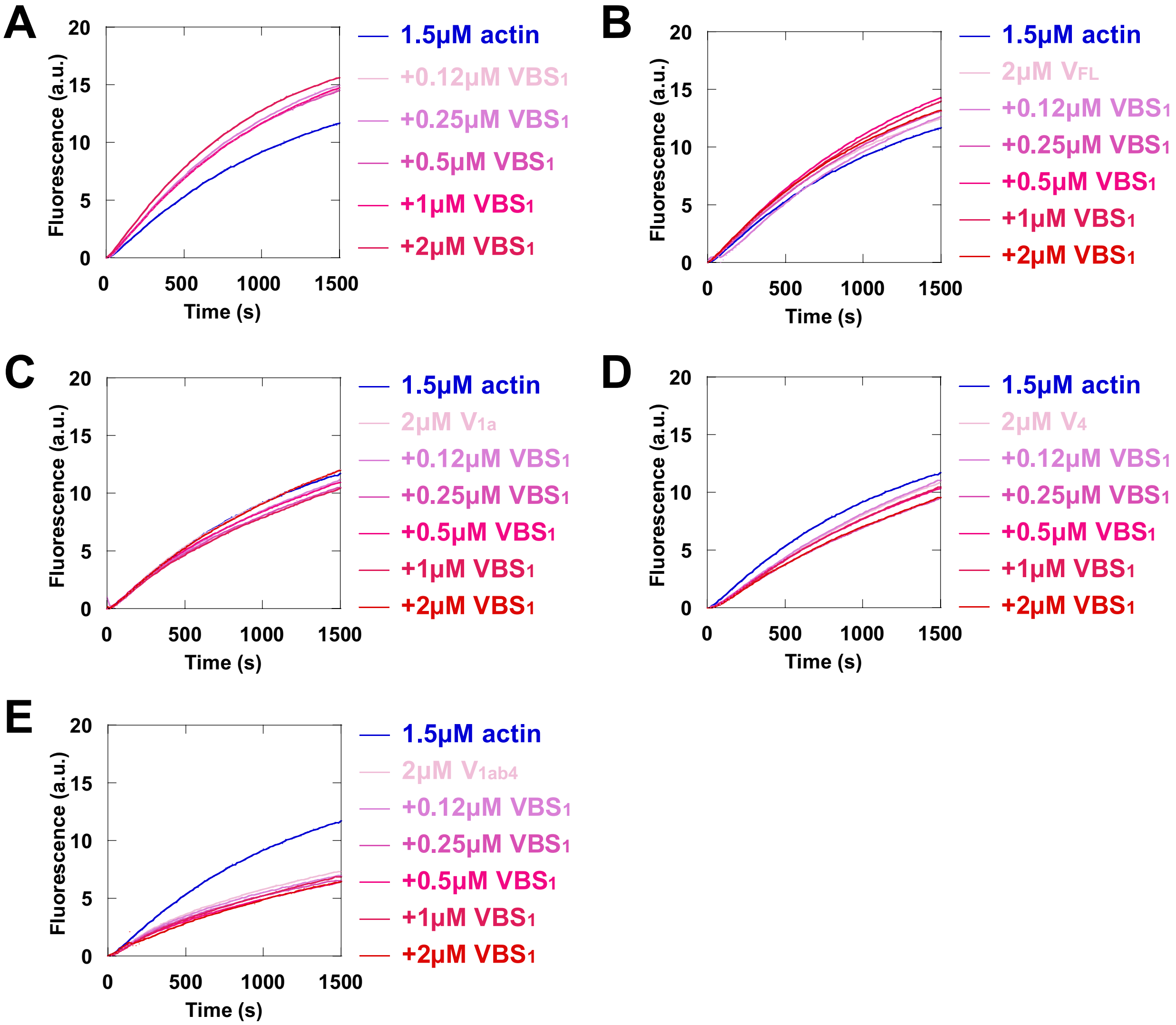


**Supplementary figure 7. Talin $VBS_1$ combined with D4-$V_t$ release has no effect on barbed-end capping by vinculin. (A-E)** The elongation of actin filament barbed end was measured in the presence of 2 μM of the indicated vinculin mutants with increasing concentrations of $VBS_1$ (0.12, 0.25, 0.5, 1, 2 μM), 100 pM spectrin-actin seeds and 1.5 μM actin (10% pyrenyl-labeled). The control kinetics showing the polymerization of 1.5 μM actin alone are the same in all panels.

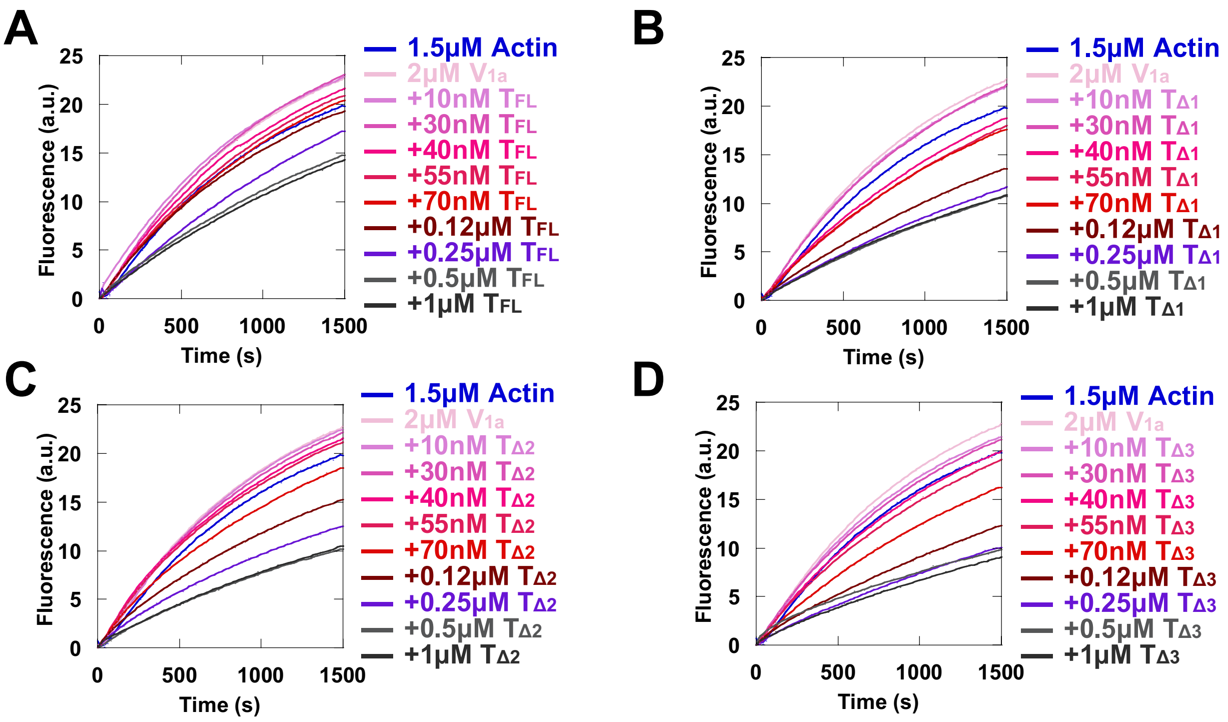


**Supplementary figure 8. A complex composed of full-length talin with exposed VBSs and V$_{1a}$ caps actin filament barbed ends. (A-D)** The elongation of actin filament barbed end was measured in the presence of 2 μM V$_{1a}$ with increasing concentrations of the indicated talin mutants, 100 pM spectrin-actin seeds and 1.5 μM actin (10% pyrenyl-labeled). The control kinetics showing the polymerization of 1.5 μM actin in the presence of 2 μM V$_{1a}$ are the same in all panels.

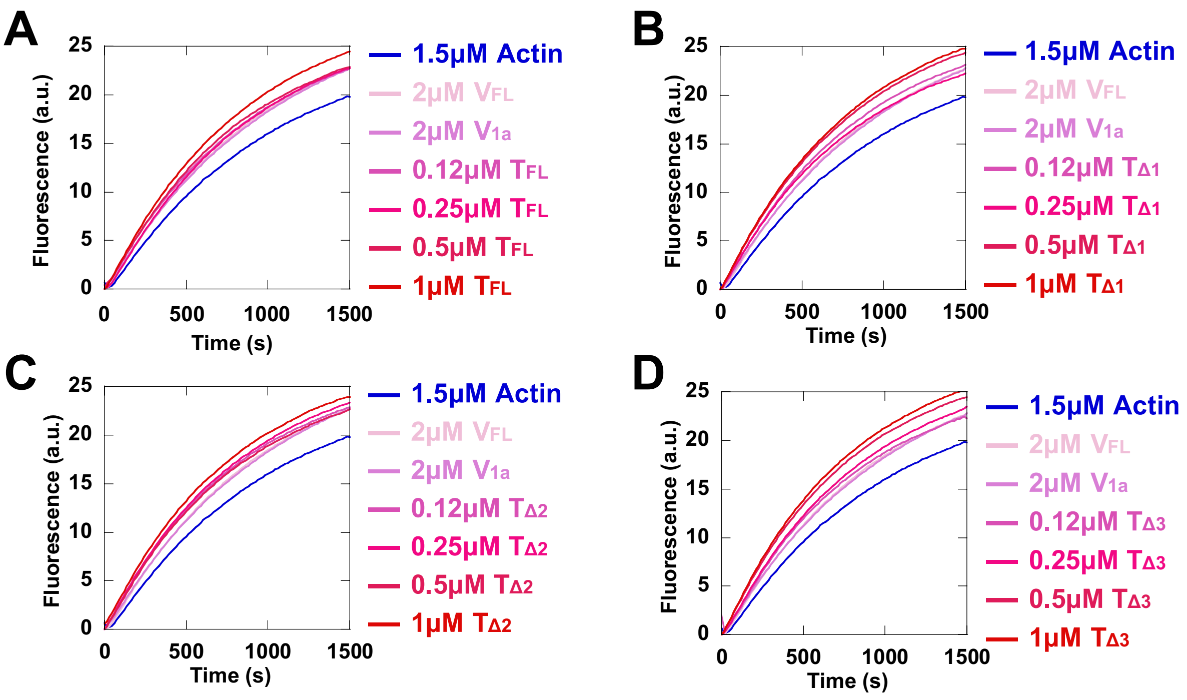


**Supplementary figure 9. Talin mutants with exposed VBSs alone do not cap barbed ends. (A-D)** The elongation of actin filament barbed end was measured in the presence of increasing concentrations of the indicated talin mutants, 100 pM spectrin-actin seeds, 1.5 μM actin (10% pyrenyl-labeled). The kinetics with 1.5 μM G-actin, 2 μM $V_{FL}$ or 2 μM $V_{1a}$, used here as additional negative controls, are the same in all panels.

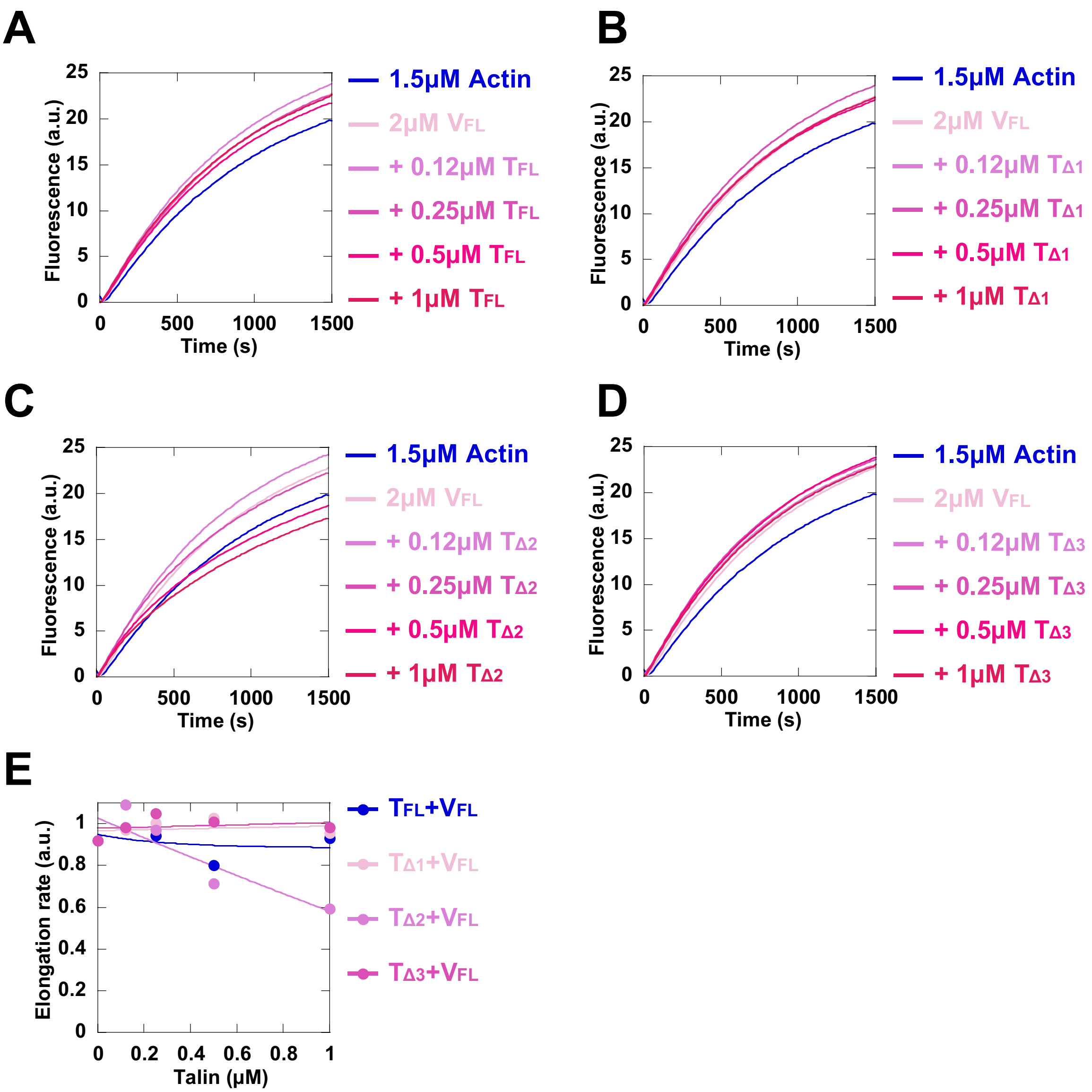


**Supplementary figure 10. Addition of full-length talin with exposed VBSs to $V_{FL}$ does not cap actin filament barbed ends. (A-D)** The elongation of actin filament barbed end was measured in the presence of 2 µM $V_{FL}$, increasing concentrations of the indicated talin mutants, 100 pM spectrin-actin seeds, 1.5 µM actin (10% pyrenyl-labeled). The kinetics with 1.5 µM actin and 2 µM $V_{FL}$ are the same in all panels. **(E)** The fraction of barbed end elongation was calculated as the ratio of the elongation rate in the presence of $V_{FL}$ and talin mutants to the elongation rate of 1.5 µM actin alone.

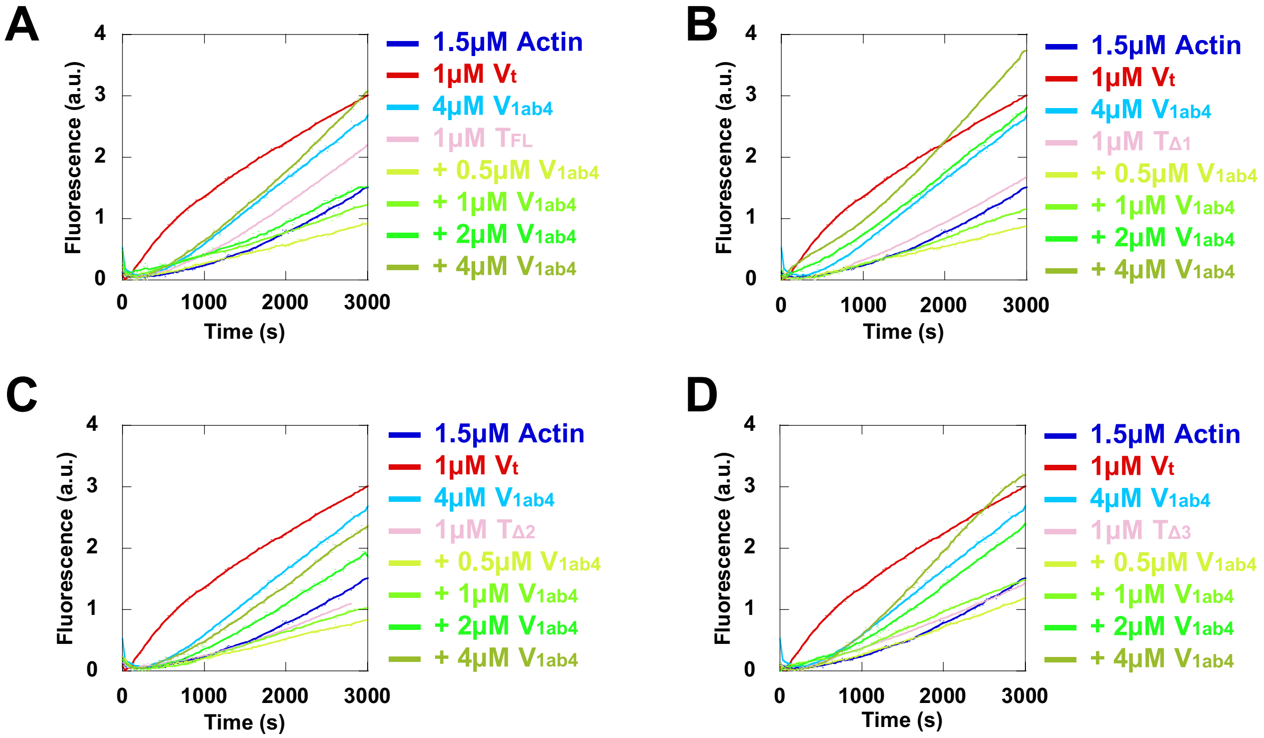


**Supplementary figure 11. Talin mutants with exposed VBSs combined with $V_{1ab4}$ do not stimulate actin assembly. (A-D)** Spontaneous actin polymerization was measured in the presence of 1μM of the indicated talin mutants, increasing concentrations of $V_{1ab4}$ and 1.5 μM actin (10% pyrenyl-labeled) in a low salt buffer (25 mM KCl). The kinetics with 1.5 μM actin and 1 μM $V_t$ alone, used as a positive control for nucleation, or 1.5 μM actin and 4 μM $V_{1ab4}$ alone are the same in all panels.

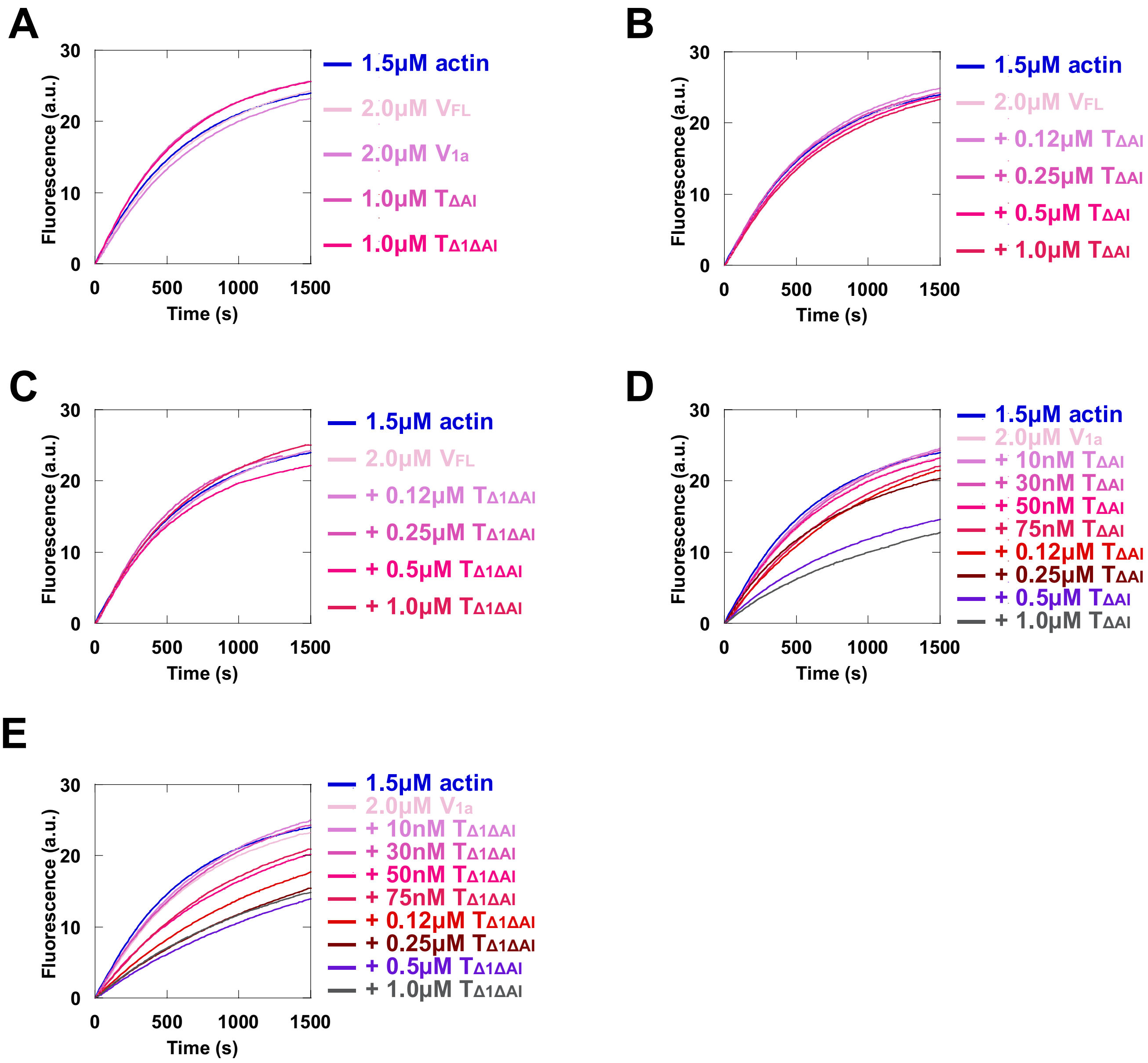


**Supplementary figure 12. Talin mutants, with released autoinhibitory contacts and one exposed VBS combine with $V_{1a}$ to cap actin filament barbed ends.** The elongation of actin filament barbed end was measured in the absence and presence of 2 µM $V_{FL}$ or 2 µM $V_{1a}$ or 1 µM $T_{\Delta AI}$ or 1 µM $T_{\Delta 1\Delta AI}$ **(A)**, in the presence of increasing concentrations of indicated talin mutants and 2 µM $V_{FL}$ **(B-C)** or 2 µM $V_{1a}$ **(D-E)**, 100 pM spectrin-actin seeds and 1.5 µM actin (10% pyrenyl-labeled). The control kinetics with 1.5 µM actin and 2 µM $V_{FL}$ **(A-C)** or 1.5 µM actin and 2 µM $V_{1a}$ **(D-E)** are the same in all panels.

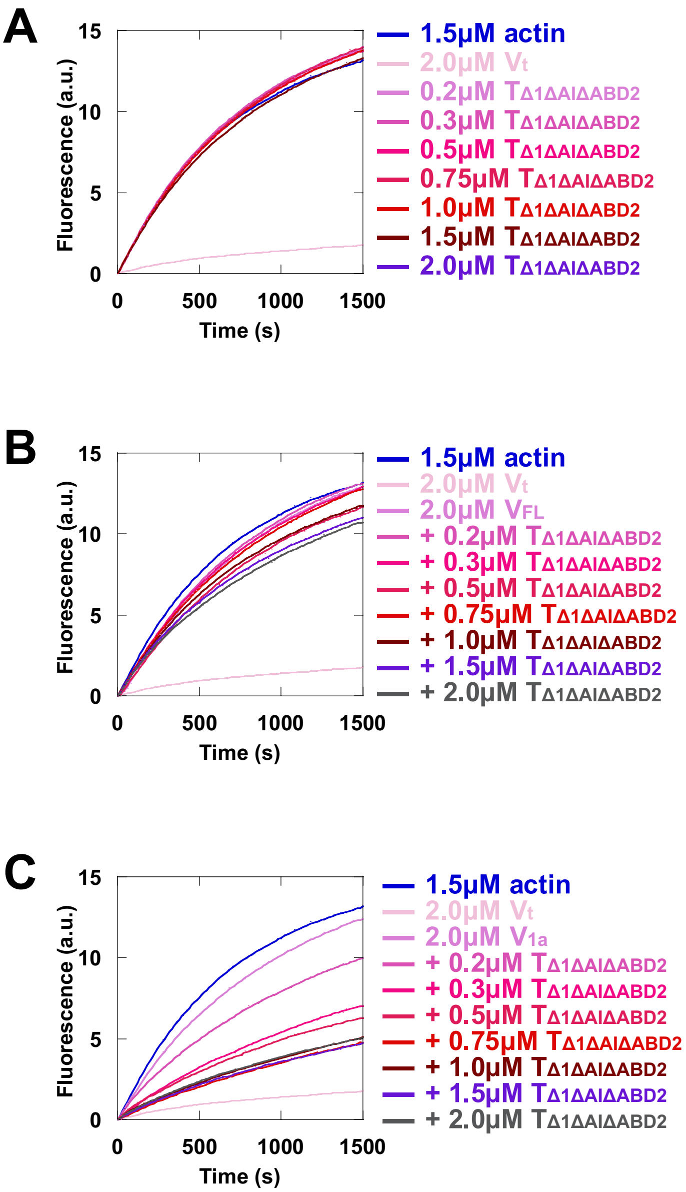


**Supplementary figure 13. A talin mutant, with released autoinhibitory contacts, one exposed VBS and lacking ABD2, combines efficiently with $V_{1a}$, but not with $V_{FL}$, to cap actin filament barbed ends. (A-C)** The elongation of actin filament barbed end was measured in the presence of increasing concentrations of $T_{\Delta1\Delta AI\Delta ABD2}$, 100 pM spectrin-actin seeds and 1.5 μM actin (10% pyrenyl-labeled) (A), increasing concentrations of $T_{\Delta1\Delta AI\Delta ABD2}$, 2 μM $V_{FL}$ (B) or 2 μM $V_{1a}$ (C), 100 pM spectrin-actin seeds and 1.5 μM actin (10% pyrenyl-labeled). The kinetics with 1.5 μM actin alone and 1.5 μM actin and $V_t$ are the same in all panels.

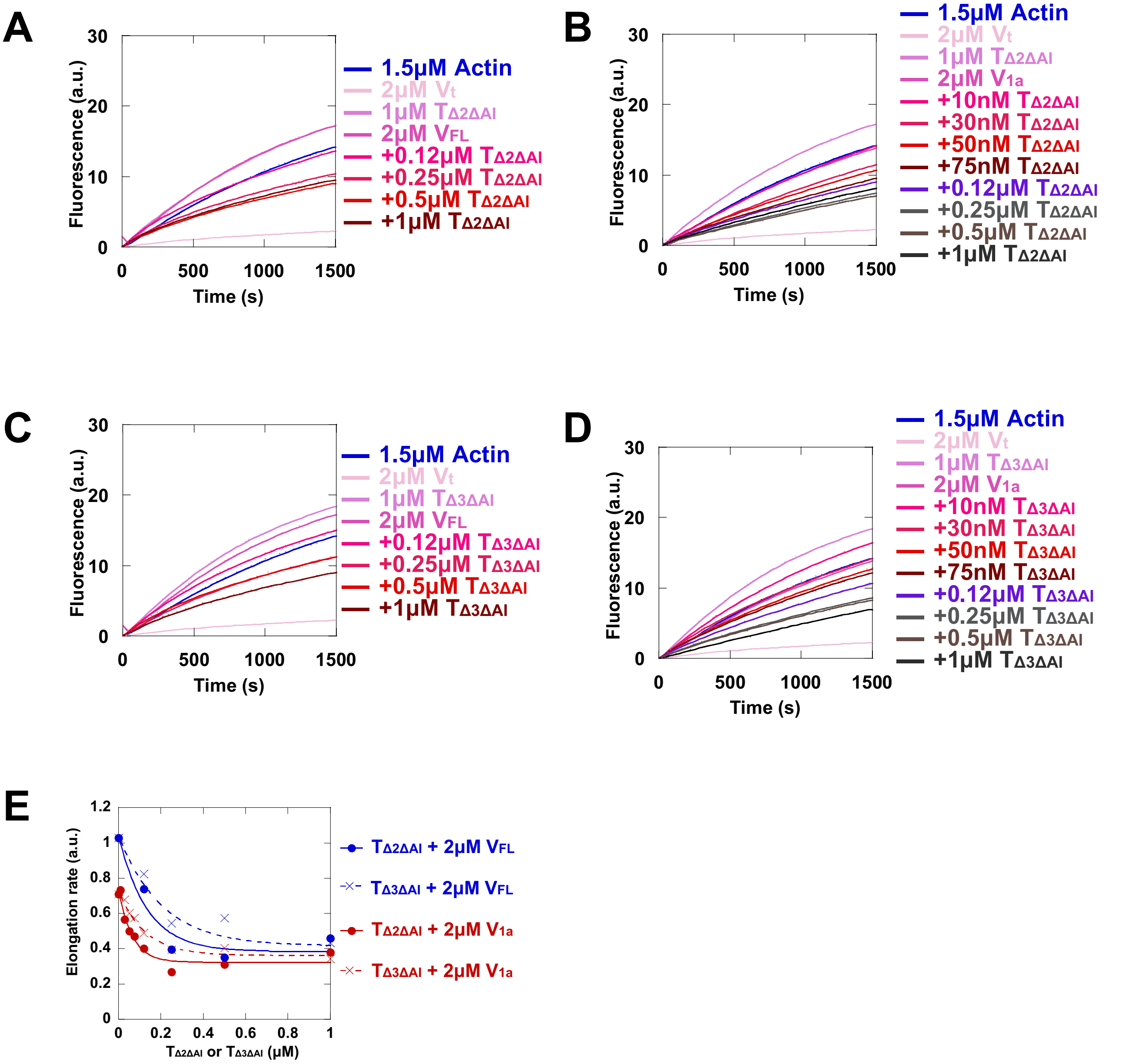


**Supplementary figure 14. Talin mutants, with released autoinhibitory contacts and two or three exposed VBSs, combine with $V_{FL}$ and $V_{1a}$ to cap actin filament barbed ends. (A-E)** The elongation of actin filament barbed end was measured in the presence of increasing concentrations of $T_{\Delta2\Delta AI}$ and 2 μM of $V_{FL}$ **(A)**, increasing concentrations of $T_{\Delta2\Delta AI}$ and 2 μM of $V_{1a}$ **(B)**, increasing concentrations of $T_{\Delta3\Delta AI}$ and 2 μM of $V_{FL}$ **(C)**, increasing concentrations of $T_{\Delta3\Delta AI}$ and 2 μM of $V_{1a}$ **(D)**, 100 pM spectrin-actin seeds and 1.5 μM actin (10% pyrenyl-labeled). The control kinetics with 1.5 μM actin and 2 μM $V_t$ or 1 μM $T_{\Delta2\Delta AI}$ or 1 μM $T_{\Delta3\Delta AI}$ or 2 μM $V_{FL}$ or 2 μM $V_{1a}$ are the same in all panels. **(E)** Quantification of the experiments in (A-D).

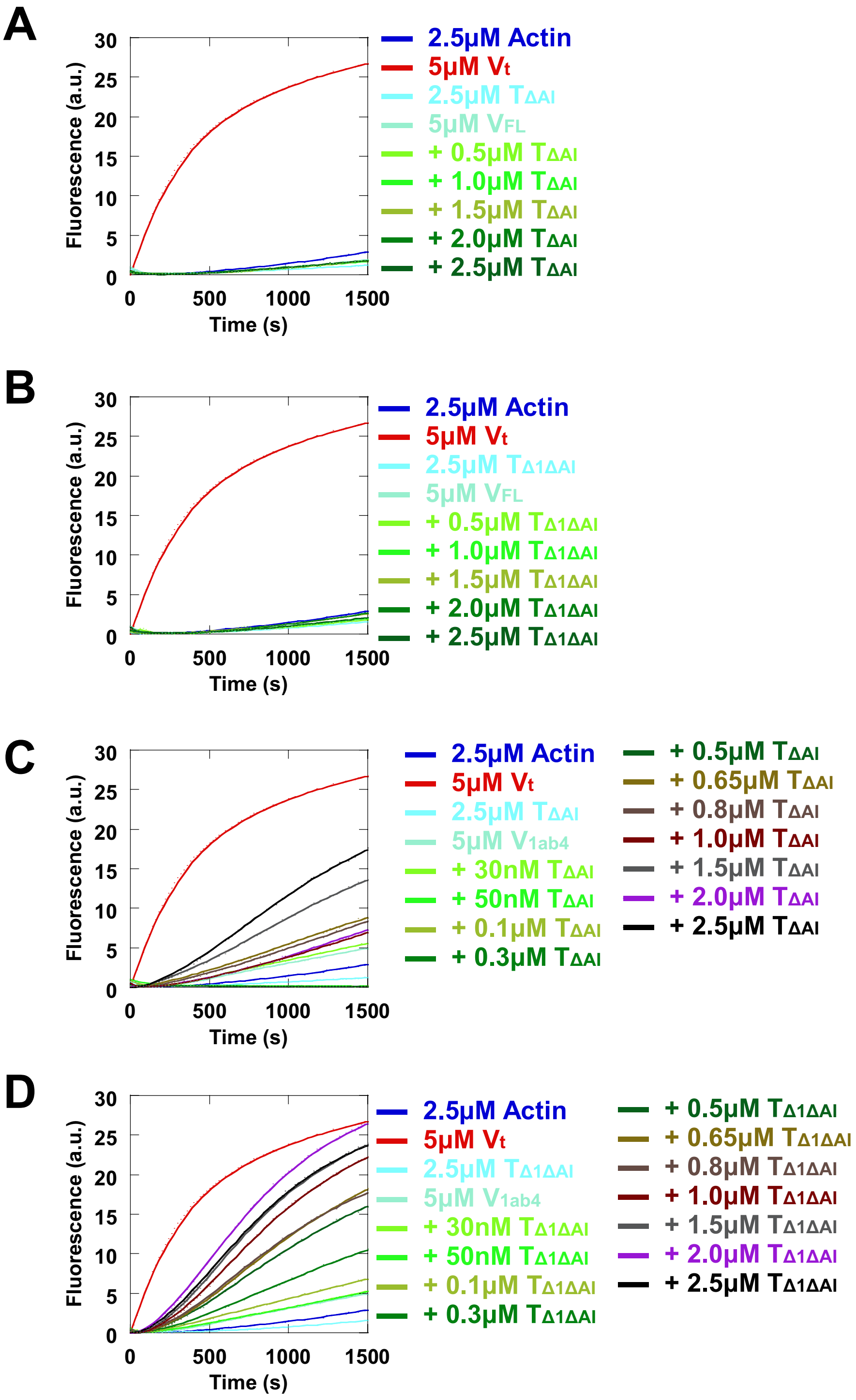


**Supplementary figure 15. Talin mutants, with released autoinhibitory contacts and one exposed VBS, combine efficiently with $V_{1ab4}$ to stimulate actin nucleation. (A-D)** Actin polymerization was measured in the absence and presence of 5 µM of $V_{FL}$ or $V_{1ab4}$ with increasing concentrations of $T_{\Delta AI}$ or $T_{\Delta 1\Delta AI}$ and 2.5 µM G-actin (10% pyreyl-labeled) in a low salt buffer (25 mM KCl). The kinetics with 2.5 µM actin alone, used as a negative control, and 2.5 µM actin and 5 µM $V_t$, used as a positive control for nucleation, are the same in all panels.

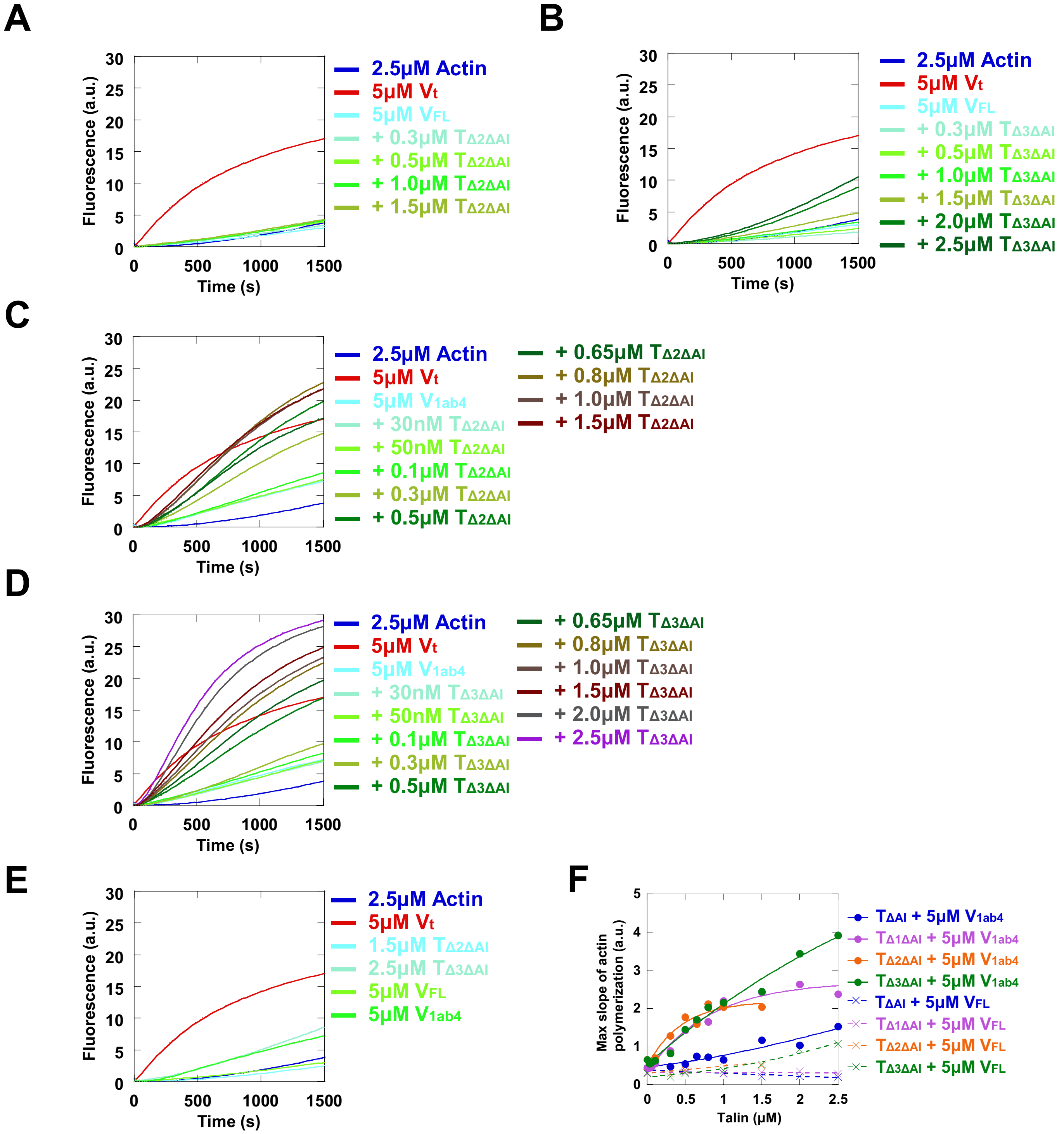


**Supplementary figure 16. Talin mutants, with released autoinhibitory contacts and two or three exposed VBSs, combine efficiently with $V_{1ab4}$ to stimulate actin nucleation. (A-D)** Actin polymerization was measured in the presence of 5 μM of $V_{FL}$ or $V_{1ab4}$ with increasing concentrations of $T_{\Delta2\Delta AI}$ or $T_{\Delta3\Delta AI}$ and 2.5 μM actin (10% pyreyl-labeled) in a low salt buffer (25 mM KCl). **(E)** Additional negative control showing the polymerization of 2.5 μM actin (10% pyreyl-labeled) in the presence of 5 μM $V_{FL}$ or 5 μM $V_{1ab4}$ or 1.5 μM $T_{\Delta2\Delta AI}$ or 2.5 μM $T_{\Delta3\Delta AI}$. The kinetics with 2.5 μM actin alone, used as a negative control, and 2.5 μM actin and 5 μM $V_t$, used as a positive control for nucleation, are the same in all panels. **(F)** The maximal rate of spontaneous actin polymerization is plotted against increasing concentrations of talin mutants in the presence of $V_{FL}$ or $V_{1ab4}$ as described in (A-D). Data for $T_{\Delta1\Delta AI}$ + $V_{FL}$ or $V_{1ab4}$ and $T_{\Delta AI}$ + $V_{FL}$ or $V_{1ab4}$ are the same as in Figure 3D.

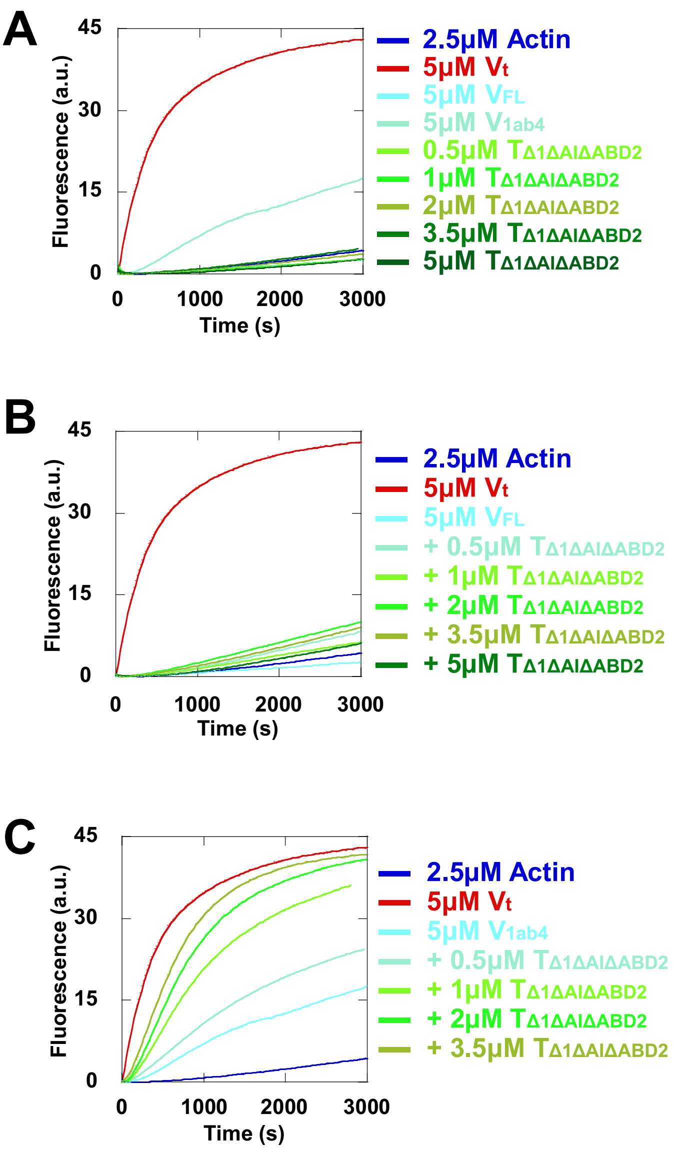


**Supplementary figure 17. A talin mutant, with released autoinhibitory contacts, one exposed VBS and lacking ABD2, combines efficiently with $V_{1ab4}$ to stimulate actin nucleation. (A-C)** Spontaneous actin polymerization was measured in the presence of 2.5 µM actin (10% pyreyl-labeled) and increasing concentrations of $T_{\Delta1\Delta AI\Delta ABD2}$ alone (A), increasing concentrations of $T_{\Delta1\Delta AI\Delta ABD2}$ and 5 µM $V_{FL}$ (B) and increasing concentrations of $T_{\Delta1\Delta AI\Delta ABD2}$ and 5 µM $V_{1ab4}$ (C) in a low salt buffer (25 mM KCl). The kinetics with 2.5 µM actin and 5 µM $V_t$ alone, used as a positive control or 5 µM $V_{1ab4}$ alone are the same in all panels.

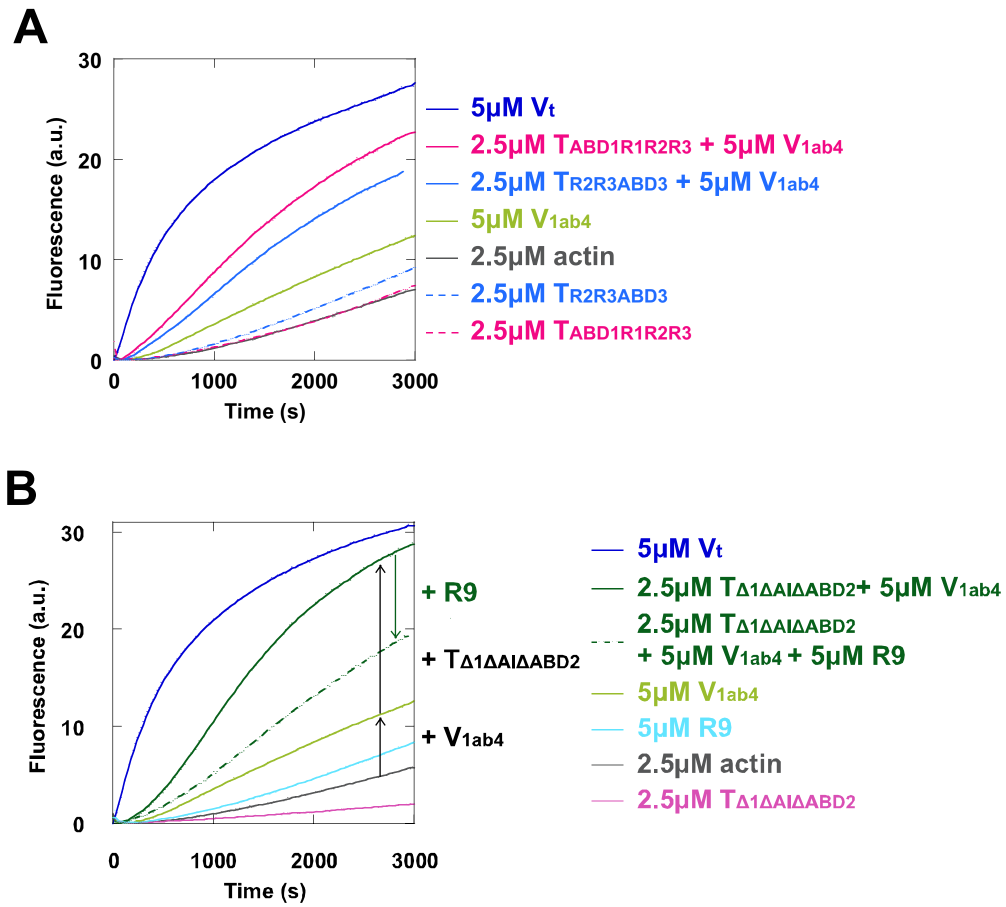


**Supplementary figure 18. Importance of talin ABDs for the activities of the talin-vinculin complex. (A,B)** Spontaneous actin polymerization was measured in presence of 2.5 μM actin (10% pyrenyl-labeled) and the indicated talin and vinculin mutants in a low salt buffer (25 mM KCl). The black arrows in (B) indicate the stimulation of actin polymerization by $V_{1ab4}$ alone and $V_{1ab4}$ + $T_{\Delta1\Delta AI\Delta ABD2}$. The dark green arrow in (B) indicates that R9 reverses the actin polymerization induced by $V_{1ab4}$ + $T_{\Delta1\Delta AI\Delta ABD2}$.

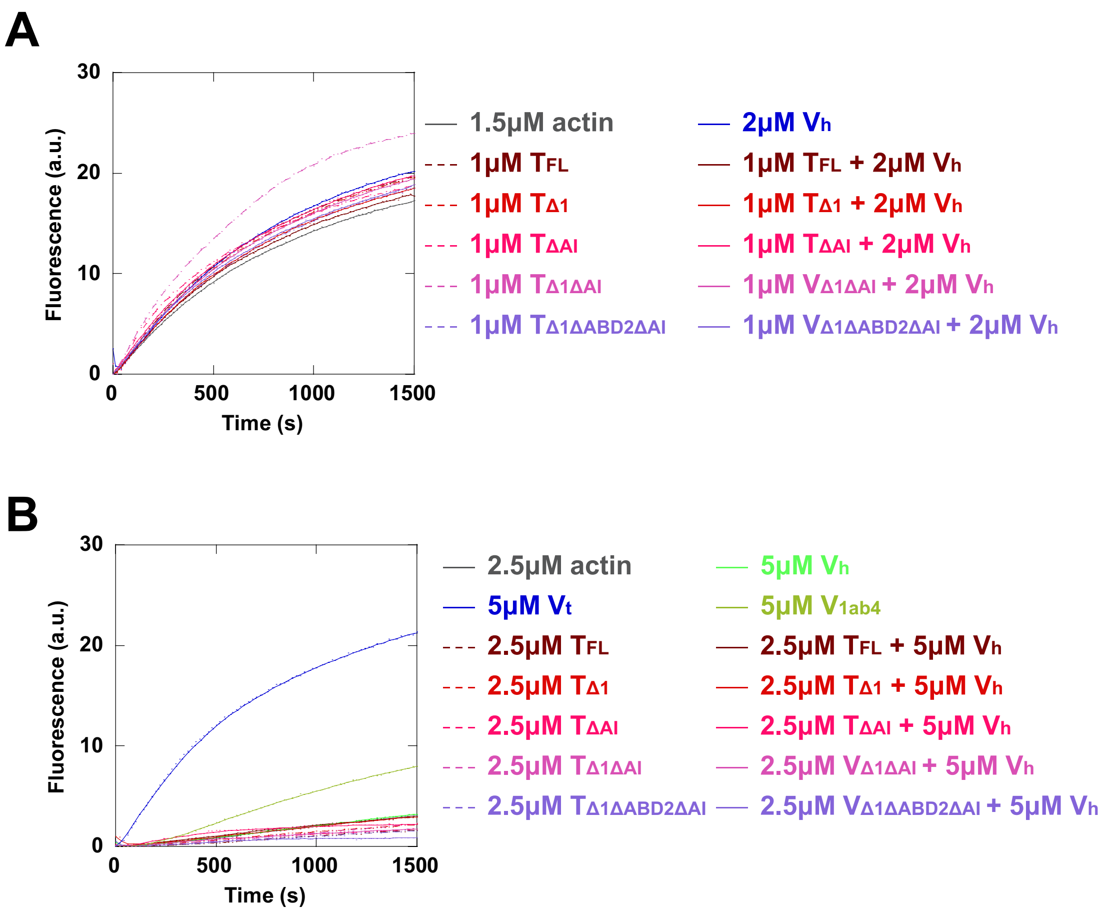


**Supplementary figure 19. A talin-vinculin complex, in which vinculin lacks $V_t$, does not nucleate and cap actin filaments. (A)** The elongation of actin filament barbed end was measured in the presence of the indicated proteins, 100 pM spectrin-actin seeds, 1.5 μM actin (10% pyrenyl-labeled). **(B)** Spontaneous actin polymerization was measured in presence of 2.5 μM actin (10% pyrenyl-labeled) and the indicated proteins in a low salt buffer (25 mM KCl).
